# The Role of Hvem and its Interaction with Btla and Cd160 in b-Cell Lymphoma Progression

**DOI:** 10.1101/754291

**Authors:** Carla Yago-Diez de Juan

## Abstract

Despite the fact that the cell surface receptor HVEM (TNFRSF14) appears to be implicated in the development and progression of B-cell lymphomas, its specific role in these tumours is still unclear. On the one hand, HVEM over-expression is related to worse prognosis in some types of B-cell lymphoma and other solid tumours. On the other hand, most mutations of HVEM in B-cell lymphomas are thought to promote tumour growth through the loss of function. Here, we used a CRISPR-Cas9 system to study the effect of HVEM loss on gene expression in a murine model of A20 B-cell lymphoma (belonging to the diffuse large B-cell lymphoma group). We show that loss of HVEM does not affect the doubling rate of A20 tumour cells in culture, but leads to a decrease in BTLA expression. HVEM-deficient A20 cells do not present a different pattern of metastatic dissemination to lymphoid organs compared with unmodified A20 cells. However, we observed a significant expansion of endogenous B-cells as a result of A20 tumour implantation in the thymus. Although we found no differences in the dissemination or progression of HVEM-deficient A20 cells, our results reveal that loss of HVEM alters the leukocyte recruitment capacity of A20 cells in hepatic tumour nodules at the intermediate stage of tumour development, which may be of relevance as a mechanism of immune evasion.

## INTRODUCTION

HVEM (Herpes Virus Entry Mediator; TNFRSF14) is a cell surface receptor which seems to play a critical role in the progression and development of B-cell lymphomas (Pasero et al. 2012; Launay et al. 2012; Upadhyay et al. 2015; Lohr et al. 2012; M’Hidi et al. 2009; Ward-Kavanagh et al. 2016). HVEM is expressed in many cell types, both hematopoietic (B and T lymphocytes, immature dendritic cells, natural killer (NK) cells, regulatory T cells (Tregs), etc.) and non-hematopoietic (stromal) cells (Pasero & Olive 2013). HVEM presents as conventional TNFRSF ligands, LIGHT (lymphotoxin-like that exhibits inducible expression) and lymphotoxin-alfa (LT-α). Mainly, it binds ligands of the immunoglobulin (Ig) superfamily, B and T lymphocyte attenuator (BTLA) and CD160, as well as glycoprotein-D (gD) of herpes simplex virus (HSV) (Steinberg et al. 2011).

In homeostatic conditions, HVEM and BTLA are usually expressed in B and T lymphocytes interacting in the cis configuration (i.e. on the same cell surface) (Murphy et al. 2006; Steinberg et al. 2011). The cis HVEM-BTLA complex induces the inhibitory signalling through the receptor BTLA, preventing effector functions and activation of survival and proliferation gene expression mediated by NF-κB transcription factor (Steinberg et al. 2011; Shui et al. 2011; Croft 2005). In addition, the cis HVEM-BTLA interaction blocks the HVEM engagement in trans (i.e. between adjacent cells) with BTLA (or CD160) (Pasero et al. 2012; Ware & Sedy 2011). After immune response activation, the stoichiometric balance of HVEM and BTLA expression is broken, and HVEM expression levels decrease while BTLA expression increases (Murphy et al. 2006). In addition, the LIGHT expression is induced on activated T lymphocytes Murphy et al. 2006). This leads to the break of the stable cis HVEM-BTLA complex, allowing the LIGHT-HVEM binding that promotes bi-directional co-stimulation of immune effector responses (Steinberg et al. 2011; Ware & Sedy 2011). It should be noted that the HVEM receptor engagement by BTLA/CD160 or LIGHT activates NF-κB transcription factor, stimulating cell proliferation and survival (Ware & Sedy 2011).

Recent findings have revealed the importance of HVEM-BTLA interaction in the B-cell lymphoma progression and prognosis (Boice et al. 2016; Pasero & Olive 2013; Ware & Sedy 2011; Gertner-Dardenne et al. 2013), although its function in the prognosis of this disease is very controversial. On the one hand, in some B-cell lymphomas, such as B-chronic lymphocytic leukemia (B-CLL), HVEM is expressed in high levels and it is related to lymphoma promotion and development (Pasero & Olive 2013). This association is supported by studies of other tumour types, such as sporadic breast cancer, melanoma, ovarian and colorectal cancer, where HVEM mutations are associated with its over-expression, which contributes to tumour progression and poor prognosis (Li et al. 2013; Inoue et al. 2015; Zhang et al. 2016). In these cases, HVEM is used by tumour cells as a ligand of the co-inhibitory BTLA receptor expressed on tumour-specific cytotoxic CD8 T cells, limiting their proliferation and survival capacity and inhibiting their cytotoxic effector activity against the tumour (Figure 1A) (Wang et al. 2005; Upadhyay et al. 2015; Gertner-Dardenne et al. 2013; Zhang et al. 2016). These results were supported by studies with HVEM knock-out (HVEM^−/−^) mice, in which HVEM^−/−^ T cells exhibit an exacerbated immune response (Wang et al. 2005). In addition, the blockade of the HVEM-BTLA interaction by means of antagonist anti-BTLA antibodies produces an increase in the number of T cells co-cultured with HVEM positive B-cell lymphoma (Gertner-Dardenne et al. 2013). In these cases, expression of HVEM could be interpreted as an immune evasion mechanism that would translate in enhanced tumour progression.

**Figure 1.**
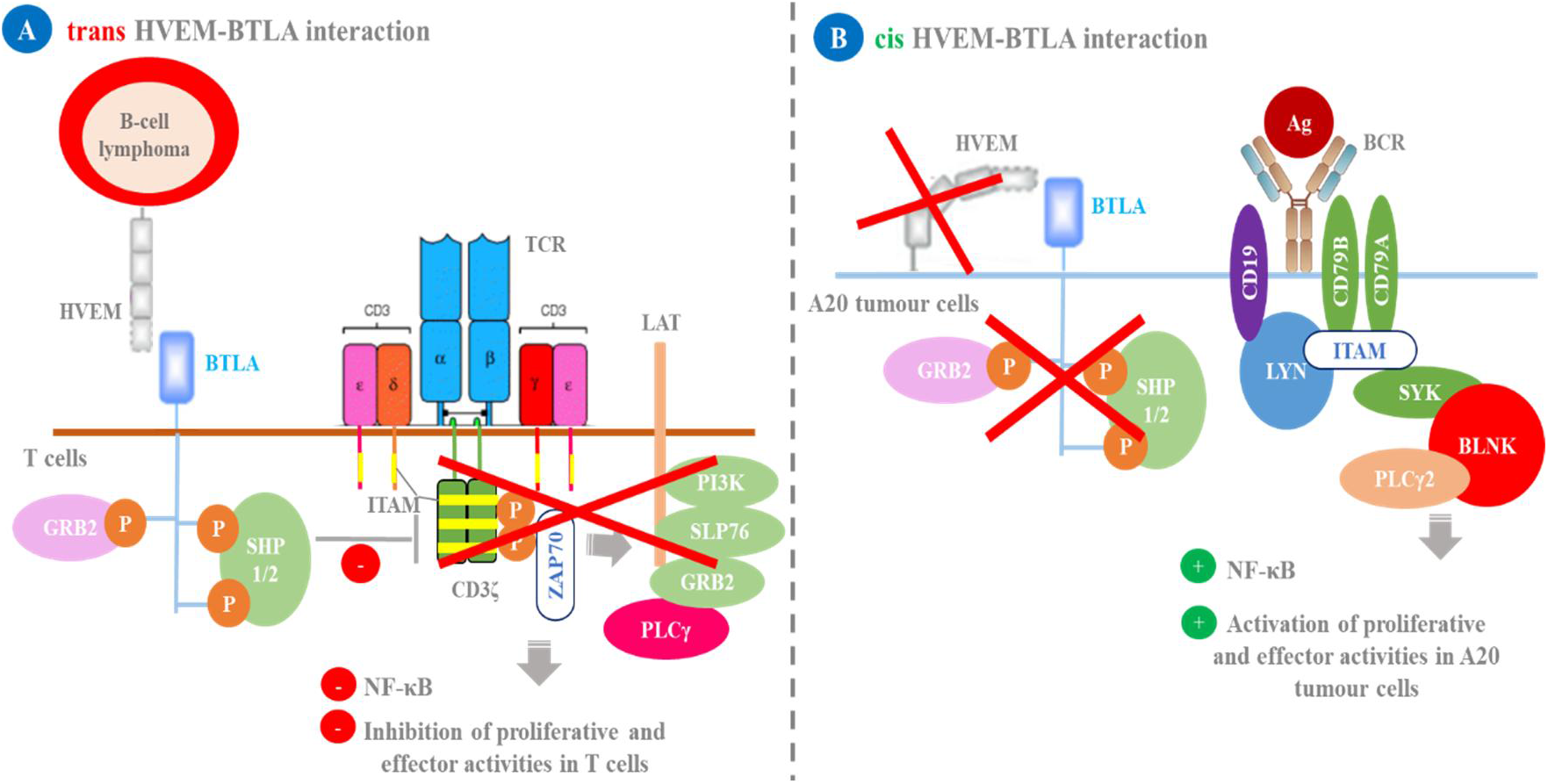
Two possible roles of HVEM in B-cell lymphoma development and progression. The trans-interaction of HVEM expressed in B-cell lymphoma, and BTLA, expressed in T cells, leads to the inhibition of TCR-mediated signalling in T cells (Wu et al. 2007). The activation of BTLA, after binding of HVEM, leads to dephosphorylation of tyrosine-based activation motifs (ITAMs) of the CD3ζ complex of TCR, preventing the activation of the protein tyrosine kinase ZAP70. In this way, ZAP70 is unable to phosphorylate and activate LAT (activation linker in T cells), which prevents the activation of a series of proteins (PI3K, SLP76, GRB2 and PLCγ). All this results in the inhibition of NF-κB activation, limiting the proliferation and responses mediated by T cells (Lieping Chen and Dallas B. Flies 2013). (B) The loss of HVEM expression in B-cell lymphoma as a mechanism for promoting tumour proliferation and development. If HVEM is not expressed on the tumour cell surface, it cannot interact in cis with BTLA. The latter is thus unable to activate its inhibitory-signalling cascade, which allows BCR-mediated signalling, activating a series of proteins such as Syk, which is phosphorylated by Lyn. Syk phosphorylates BLNK and phospholipase Cγ2 (PLCγ2), finally activating NF-κB factor and the proliferation and effector responses mediated by B-cells (Boice et al. 2016).

On the other hand, in the two most common subtypes of B-cell lymphoma, follicular lymphoma (FL) and diffuse large B-cell lymphoma (DLBCL), the majority of *TNFRSF14* gene mutations have been associated with the loss of HVEM expression on the cell surface, favouring tumour development and progression (Boice et al. 2016; Lohr et al. 2012; Cheung et al. 2010; Launay et al. 2012; Pasqualucci & Dalla-Favera 2018). The loss of HVEM expression in FL prevents the cis HVEM-BTLA interaction, which results in a decrease in the activation threshold of tumour B-cells (Boice et al. 2016) (Figure 1B). Therefore, these tumour cells are more predisposed to uncontrolled proliferation. Furthermore, it has been shown that HVEM deficient FL promotes a supportive microenvironment to tumour development by the recruitment of follicular helper cells (T_FH_) (Boice et al. 2016). Hence, HVEM acts as a tumour suppressor in this context.

It is clear that HVEM has a role in B-cell lymphoma development and progression, but there is disagreement about whether its function promotes or prevents tumour development and progression. Here, we investigated whether suppression of HVEM expression by CRISPR-Cas9 editing in an A20 B-cell lymphoma (belonging to the DBCL group) (Kim et al. 1979), affects tumour development and progression in an experimental murine model of B-cell lymphoma.

## MATERIAL AND METHODS

### Mice

In order to generate a murine model of A20 B-cell lymphoma were used 6-8 weeks-old female syngeneic Balb/c (H-2^d^) mice (Kim et al. 1979). These mice were inoculated intravenously in the tail with the different A20 tumour cell lines to evaluate in vivo tumour development and progression. Animals were bred at the animal facility of the University of Leon (Spain) in standard light/darkness conditions and free-accede to water and meal. All experiments with animals had been made in accordance with the Ethical Committee for Animal Research of the School of Veterinary Medicine (University of Leon) and the European Guidelines for Animal Care and Use of Laboratory Animals.

### Mammalian cell lines and growth conditions

A20 B-cell lymphoma line was obtained spontaneously from a reticular cell of Balb/cAnN mouse in advanced stage (Passineau 2005) (Material Transfer Agreement (MTA), Dr Arnab Ghosh, New York). HEK 293T cells were obtained from American Type Culture Collection (ATCC). Hepa 1-6 cells were derived from mouse liver carcinoma (MTA, Dr I. Anegon, Nantes) (Darlington et al. 1980). Cells were cultured in Iscove’s Modified Dulbecco’s medium (Gibco) supplement with 10% fetal bovine serum (FBS) (Gibco), or in RPMI 1640 medium (Gibco) supplemented with 7% FBS, 2 mM L^−^ glutamine, 1 mM sodium pyruvate, 10 mM HEPES, 50 μg/mL gentamicin and 5 × 10^−5^ M β-mercaptoethanol (Sigma-Aldrich). Cells were incubated at 37°C in 5% atmospheric CO_2_, and they were split every three days to prevent them from becoming completely confluent. 0.05% Trypsin-EDTA was used in order to split adherent cells.

### Transduction of A20 B-cell lymphoma with lentivirus expressing eGFP or Cas9 nuclease

We used lentivirus expressing enhanced green fluorescent (eGFP) protein (given by the Center for Energy, Environmental and Technological Research (CIEMAT), Spain) or *Streptococcus pyogenes* Cas9 (spCas9) nuclease, in order to generate A20 cells with stable GFP marking or Cas9 expression, respectively. To obtain the lentiviral vectors encoding for Cas9 were used the following plasmids: (i) spCas9-PURO (D. Heckel et al. 2014) (MTA) plasmid expressing Cas9 nuclease and puromycin resistance, (ii) pPAX2, a second-generation lentiviral packaging plasmid (Dr. P. Schneider, MTA), and (iii) pVSV-g plasmid expressing lentiviral envelope (G glycoprotein of the vesicular stomatitis virus) (Dr. P. Schneider, MTA) (Stewart et al. 2003). To pack and produce the lentiviral particles were used HEK293T cells. Besides, due to these A20 tumour cells present a high infection multiplicity (MOI) (50-100) (Serafini et al. 2004; Anastasov et al. 2009), the desirable viral titer (≥ 10^7^ transduction units (TU)) was achieved by ultracentrifugation (20.000 rpm, 2h at 4°). The A20 cells infection was made in presence of polybrene (8 μg/mL) (Davis et al. 2002) and retronectin (Clontech), which facilitates the co-localization between tumour cells and lentiviral particles, improving the process of transduction (Kitamura et al. 2003). Furthermore, given that A20 tumour cells grow in suspension, the infection process was complemented by spinoculation, which is centrifugation at 2400 rpm for 1 h 30 min at room temperature (RT) (Ye et al. 2008). Spinoculation increases the contact between lentiviral particles and A20 suspension cells, which enhances the infection efficiency.

On the one hand, to obtain A20 tumour cell line with stable GFP expression, A20 tumour cells transduced with GFP lentiviral particles were cloned by limited dilution in 96-well plates. First, we made a previous A20 GFP positive clones selection by fluorescence microscopy, and then we corroborated GFP expression by flow cytometry and selected A20 GFP clone 2D9.

On the other hand, to generate A20 tumour cell line with stable Cas9 nuclease expression. A20 GFP cells transduced with Cas9 lentiviral particles were selected with puromycin (1.5 μg/mL) (Thermo Scientific) for 15 days. After the puromycin selection, A20 GFP cells resistant to the treatment were cloned by limited dilution in 96-well plates. Clones were analysed by PCR for Cas9 amplification using these primers: forward, 5’-CTGAAGAGAACCGCCAGAAG-3’; and reverse, 5’-AGCTGGGCGATCAGATTTT-3’. Finally, we selected A20 GFP Cas9 clone 1D5 with stable Cas9 integration.

### Design of sgRNAs guides sequences in order to clone them into lentiCRISPRv2 and sgRNA-tagRFP657 vectors

The different single-guide RNAs (sgRNAs) were designed from exon 1 (69 pb) of HVEM sequence (Balb/c mouse strain, GeneBank accession number: NM 019418) by *Massachusetts Institute of Technology* (MIT) software (Zhang Laboratory, 2015). This program is based on the generating of 20 pb complementary sequences to target genome zone, taking into account the presence of 5’-NGG sequence (Protospacer Adjacent Motif (PAM) sequence) in the genome.

In order to clone each sgRNA into the lentiCRISPRv2 (third-generation lentiviral plasmid coding for spCas9 nuclease, CRISPR RNA (crRNA) and transactivating CRISPR RNA (tracrRNA) complex, puromycine resistance cassette, and *BsmBI* cloning site for sgRNA) (Sanjana et al. 2015) (Addgene plasmid # 52961) and sgRNA-tagRFP657 (lentiviral plasmid with *BsmBI* cloning site for sgRNA and fluorescence protein that emits in APC (allophycocyanin) channel (coding by *RFP657* gene reporter) (D. Heckel et al. 2014) (Dr. Dirk Heckel, Hannover University, MTA) expression vectors, we included the complementary sticky ends (CACC/CAAA) resulting of the Fast Digest *BsmBI* (Esp3I) (Thermo Scientific) restriction enzyme action, and a guanine (G) in the 5’-end, which is necessary for the start of transcription (Ran et al. 2013). The oligonucleotide of each sgRNA (sgDNA) were synthesized by Sigma-Aldrich, and they were annealed to generate oligonucleotide duplex (dsDNA-guided) using this protocol: 10 μl oligo 1 (100 μM), 10 μl oligo 2 (100 μM) and 20 μl Buffer TE (10 mM Tris, 1 mM EDTA) (95°C for 4 min, 70°C for 10 min). After annealing, 5 μl dsDNA-guided (2.5 μM) was phosphorylated by 1 μl T4 polynucleotide kinase (PNK) (1U) in 5 μl T4 PNK buffer A (10x) (34 μl final volume) (Thermo Scientific).

LentiCRISPRv2 and sgRNA-tagRFP657 vectors were digested by Fast digest *BsmBI* (according to the manufacturer’s protocols). 0.5-1 μg linearized and purified plasmids were dephosphorylated by 1 μl alkaline phosphatase (1U) (FastAP) in 2 μl FastAP buffer (10x) (20 μl final volume) (Thermo Scientific) (37°C for 30 min, 70°C for 5 min).

Finally, 2 μl dsDNA-guided oligos (12.5 nM, 1/200) were cloned into linearized and dephosphorylated lentiCRISPRv2 and sgRNA-tagRFP657 plasmids (50 ng) by 1 μl T4 DNA ligase (5 U/μl) in 4 μl T4 DNA ligase buffer (5x) (Thermo Scientific) (23°C for 20 min). 21 μl ligation reaction was treated with 1 μl exonuclease Plasmid-Safe ATP-dependent (10U) in 3 μl Plasmid-Safe buffer (10x) and supplemented with 3 μl ATP (10mM) (Epicenter) (37°C for 30 min, 70°C for 30 min). This exonuclease selectively eliminates contaminating bacterial chromosomal DNA, minimizing its cloning probability (Ran et al. 2013).

Plasmid-Safe-treated product was added into ice-cold competent Stbl3 strain, which presents a low recombinant-frequency (Hanahan 1983), and incubated for 15 min. This was transformed by heat shock protocol (42°C for 45 s) and returned it immediately to the ice for 1 min. After, 400 μl SOC medium was added and the transformation was recovered on a shaking incubator at 37°C for 45 min. Finally, the transformation was seeded in plates of LB-TB with agar and carbenicillin (50 μg/mL) and incubated the plates at 37°C overnight.

### Plasmid extraction by alkaline lysis

DNA plasmid was extracted by the alkaline lysis method using *QIAprep Spin Miniprep* Kit (Qiagen) for molecular biology procedures, and the *GeneJET Plasmid Midiprep* Kit (Thermo Scientific) for transfections of eukaryotic cells, following the manufacturer’s protocol.

### Genomic DNA extraction

Genomic DNA was extracted by boiling 100.000 cells (95°C for 10 min) and using Chelex 100 resin (5% Chelex 100 buffer, 0.1% SDS, 1% NP-40, 1% Tween 20 and sterile MilliQ water). The Chelex 100 is an ion exchange resin that works by cleaning the divalent metal ions, which can inhibit processes such as PCR.

However, for procedures that require extraction of DNA with higher purity and performance, *speed tools DNA Extraction* kit (Biotools) was used with the manufacturer’s instructions.

### T7 endonuclease I assay

Hepa 1-6 cells (70-80% confluent) were transfected with 3 μg CRISPR-Cas9-sgRNA constructions (sgRNA 1, 3, 6 or 13 cloned separately into lentiCRISPRv2 vector) and 9 μg lipofectamine (Invitrogen) in Opti-MEM medium (Gibco). 5-6 days after, genomic DNA was extracted from transfected Hepa 1-6 cells to amplify HVEM target region by PCR with forward primer: 5’-GAGCTCATGGAACCTCTCCCAG-3’; or reverse primer: 5’-CTGGGAGAGGTTCCATGAGCTC-3’.

200 ng PCR product was submitted to hybridization reaction in 2 μl NEB buffer 2 and 19 μl final volume (95°C for 5 min, 95°C to 85°C at −2°C/s, 85°C to 25°C at −0.1°C/s, and finally keep at 4°C). Annealed DNA products were digested by T7 endonuclease I (10 U) (NEB, M0302) (37°C for 30 min).

Immediately afterwards, the product of the digestion was checked by electrophoresis in a 2% agarose gel. *Quantity One software* (Biorad) was used for the image analysis, and the estimated gene modification was calculated by following the formula: % *gene modification* = *100* × *(1*− *(1*-*fraction cleaved)^1/2^)*, where fraction cleaved results from the sum of cleavage product peaks divided by the sum of cleavage product and the parent peaks. (Mackay & Segal 2010).

### pRR-eGFP system

The pRR-eGFP expression vector was used to determine the cleavage efficiency of the different sgRNAs selected. The pRR-eGFP system is characterized by presenting the N-terminal (200 nt) of eGFP protein interrupted prematurely by a stop codon, followed by a multiple cloning site (MCS), and the complete coding sequence for eGFP protein preceded by another stop codon (Flemr & Bu 2015). The pRR-eGFP system is based on the re-establishment eGFP fluorescence emission through homologous recombination repair of the ends after the cleavage Cas9-mediated guided by sgRNA (Flemr & Bu 2015).

HEK 293T cells were transfected by lipofectamine with pRR-eGFP-HVEM (codifies for the extracellular domain (ECD) of HVEM) and spCas9-PURO (codifies for the nuclease Cas9) plasmid, and separately with the sgRNA-cloned-tagRFP657 plasmid (that contain each sgRNA 1, 3, 6 or 13). In addition, HEK 293T cells were transfected separately with pRR-eGFP-HVEM and pRR-eGFP plasmid to analyse the possible basal auto-fluorescence of the truncated eGFP protein. At 48 hours post-transfection, the eGFP fluorescence intensity was analysed by flow cytometry. The efficiency of HVEM edition was determined by the average intensity of fluorescence, in such a way that at higher fluorescence levels, more effective will be the sgRNA (Flemr & Bu 2015).

### Generation of a stable HVEM knock-out mutation on A20 GFP cells

For stable HVEM mutation, 1 × 10^7^ A20 GFP cells/mL in 400 μL of RPMI 1640 supplemented by 10% FBS and with 30 μg lentiCRISPRv2-sgRNA_1 were electroporated at RT in 0.4 cm cuvettes in a Gene Pulser Xcell (BioRad) at 280 V and 950 μF. Electroporated A20 GFP cells were quickly transferred to complete RPMI 1640 medium supplemented with 14% FBS (this percent is higher than usual to promote recovery of electroporated tumour cells). After 48-72h, electroporated A20 GFP cells were analysed by flow cytometer to prove if there was a population with negative HVEM expression. Finally, heterogeneous electroporated A20 GFP cells were cloned by limited dilution and > 300 clones were analysed by flow cytometry to select HVEM knock-out mutation.

### Characterization of the mutations produced by the CRISPR-Cas9 system

The mutation/s produced by the CRISPR-Cas9 system was/were characterized by sequencing of HVEM target region, amplified by PCR with forward primer: 5’-GAGCTCATGGAACCTCTCCCAG-3’; or reverse primer: 5’-CTGGGAGAGGTTCCATGAGCTC-3’ from A20 GFP HVEM^−/−^ extracted genomic DNA. The genomic DNA from unmodified A20 GFP WT cells was used as sequencing control. *Clustal bioinformatics tools* (EMBL-EBI) were used for the analysis and comparison of nucleotide and protein sequences.

### Doubling time calculation

For doubling time (DT) calculation was used the following formula: *DT = T(ln2)/ln(Xe/Xb*) (ATCC, University Blvd. Manassas, 2014), where *Xb* is the initial number of cells (2.5 × 10^4^) and *Xe* is the average of the final cell number at the determined incubation time (T). For each incubation time, three counting replicas were included for each cell type and DT of both cell lines was compared using the unpaired Student's t-test statistical analysis (considering a *p-value* < 0.05 as statistically significant).

### Off-target effects of CRISPR-Cas9 system

The first five genes with a higher off-target probability were selected to study the possible off-target effects (Supplementary Table S1). The values of each off-target sequence were calculated according to the position of the non-complementary nucleotides or mismatches (MMs) about the PAM sequence (Supplementary Table S1). In this way, the off-target sequences that present the MMs furthest from the PAM region will have higher values and will increase the off-target probability. On the other hand, it was also necessary to take into account if the off-target sequences belong to coding regions (exons) or non-coding regions (introns). Only two of off-target sequences (corresponding to *Sipa1l3* and *Oplah* genes) are located in coding regions. Off-target regions of the different genes were amplified by PCR with flanking primers from the genomic DNA extracted from A20 GFP HVEM^−/−^ and were compared with WT sequences deposited in Ensembl database using BLAST (*Basic Local Alignment Search Tool*).

### Flow cytometry

50-100 × 10^3^ cells/P96 was incubated in with FACS buffer (Hanks Balanced Salt Solution Mg^2+^ and Ca^2+^ free (1x), 0.1% bovine serum albumin (BSA) fraction V, 5 mM EDTA 2Na2H_2_O, 1% depleted FBS, 0.5% phenol red solution, 0.02% sodium azide and Milli-Q water to 1L) with primary antibody for 15-20 min, and after with secondary antibody for 8-10 min. Non-specific FcγR binding was blocked using rat anti-mouse mAb 2.4G2 (CD16/CD32). In the end, in order to exclude dead cells, 10μL/well propidium iodide (IP) was added.

To phenotypic characterization of A20 tumour cells, the following monoclonal antibodies (mAb) were used: (i) biotinylated mAbs (1 μg/mL): HVEM (6C9), BTLA (4G12b), CD160, PD1, PD-L1, and CD49d (α_4_-integrin). These biotinylated antibodies were revealed with biotinylated secondary antibodies followed by phycoerythrin-coupled streptavidin (SA-PE). (ii) Phycoerythrin (PE)-labeled mAbs (1/300): CD21/35 (7E9), CD19 (6D5), CD18 (β_2_-integrin), CD11a (α_L_-integrin), and IgM. (iii) Peridinin Chlorophyll Protein Complex (PerCP)-conjugated mAbs (1/400): IgD. As controls, IgG2a-biotinylated rat isotype (1 μg/mL), PE-isotype (1/300), and isotype-PerCP (1/400) were used.

In order to study the different immune system cells recruitment by A20 tumour in hepatic tumour nodules and tumour progression and metastasis to primary and secondary lymphoid organs, the following combination of antibodies was used: anti-CD19-PE (1/300), anti-DX5-APC (1/400), anti-TCR-β-Biotin (1μg/mL), anti-CD11b-Percp-Cy5 (1/400), anti-Ly6-C-Biotin (1μg/mL), anti-Ly6-G-PE (1/500), anti-CD3-FITC (1/300, anti-CD8-PE (1/500), anti-CD4-APC (1/400), anti-CD11c-Biotin (1μg/mL), anti-F4/80-PE (1/500). To reveal the biotinylated antibodies, streptavidin-bright Violet 421 (SA-BV421) (1/800) was used.

*CyAn ADP Flow Cytometer* (Beckman Coulter, USA) was used for flow cytometry acquisition and data were analysed with FlowJo v10 (Tree Star, Ashland, USA) and Summit v4.3 acquisition (Beckman Coulter, USA) software programs.

### Inoculation of A20 tumour cells and tumour progression monitoring in a B-cell lymphoma murine model

1 × 10^6^ A20 tumour cells (A20 GFP (2D9) or A20 GFP HVEM^−/−^ (1F1)) in 1 ml RPMI 1640 medium and 1% FBS were intravenously inoculated in the tail of Balb/c female mice from 6 to 8 weeks of age. On day 15 post-transplant, tumour progression was analysed by flow cytometry, extracting between 200-300μL blood by tail excision. For blood collection, 10 μL heparin/tube were used, preventing it from coagulating. To extract the mononuclear cells, a blood sample was subjected to a Ficoll density gradient (Thermo Scientific) by centrifugation at 1200 rpm for 20 min (without deceleration to avoid breaking the interface). The ring of mononuclear cells was isolated taking care not to drag the granulocytes/erythrocytes deposited on the bottom. It was centrifuged at 2000-3000 rpm, and the possible erythrocytes were lysed with the ACK (ammonia-chloride-potassium) lysis buffer (for 2 minutes at RT), and the reaction was stopped with FACS buffer or PBS. Subsequently, the cells were analysed by flow cytometry.

### Lymphoid organs and hepatic tumour nodules processing

Mice were sacrificed by CO_2_ inhalation at 22, 28 and 34 days post-tumour implantation. Primary and secondary lymphoid organs (thymus, bone marrow (one tibia), spleen and peripheral lymph nodes) and hepatic tumour nodules were analysed. The spleen was first mechanically processed in the ACK erythrocyte lysis buffer, and then FACS buffer was added to stop the reaction. Thymus, the peripheral lymph nodes and hepatic tumour nodules were mechanically processed in FACS buffer. Bone marrow was also treated with ACK. Resulting cells of the different organs were filtered using nylon membranes, counted and analysed by flow cytometry.

### Classic histology

Spleen, thymus, peripheral nodes, and hepatic tumour nodules were fixed with a 10% formalin solution, to be embedded in paraffin and cut into 4-5 μm sections which were stained with hematoxylin and eosin. This experimental approach was carried out with the help and collaboration of the Pathological Anatomy Unit of the Department of Animal Health (University of Leon). Finally, the preparations were analysed by optical microscopy.

### Statistical analysis

Data obtained from the experiments were analyzed using *GraphPad Prism version 6.01* program. Statistical analyses were performed applying the unpaired Student’s t-test. A p-value less than 0.05 was considered statistically significant. On the other hand, Grubb method (alpha = 0.05) was used for outliers identification and elimination.

## RESULTS

### A20 B-cell lymphoma line expresses low levels of HVEM compared with high expression of BTLA

A20 B-cell lymphoma line was analysed by flow cytometry, in order to prove the HVEM expression. Our results revealed low levels of HVEM ligand expression on the A20 cell membrane (Figure 2). Meanwhile, these A20 cells showed high-level expression of its BTLA co-inhibitory receptor (Figure 2). On the other hand, we studied CD160 and PD-1/PD-1 ligand (PD-L1) expression by A20 tumour cells. We found that they did not express the CD160 and PD-1 receptor (Figure 2). In contrast, they showed high expression of PD-L1, similar to BTLA expression (Figure 2).

**Figure 2.**
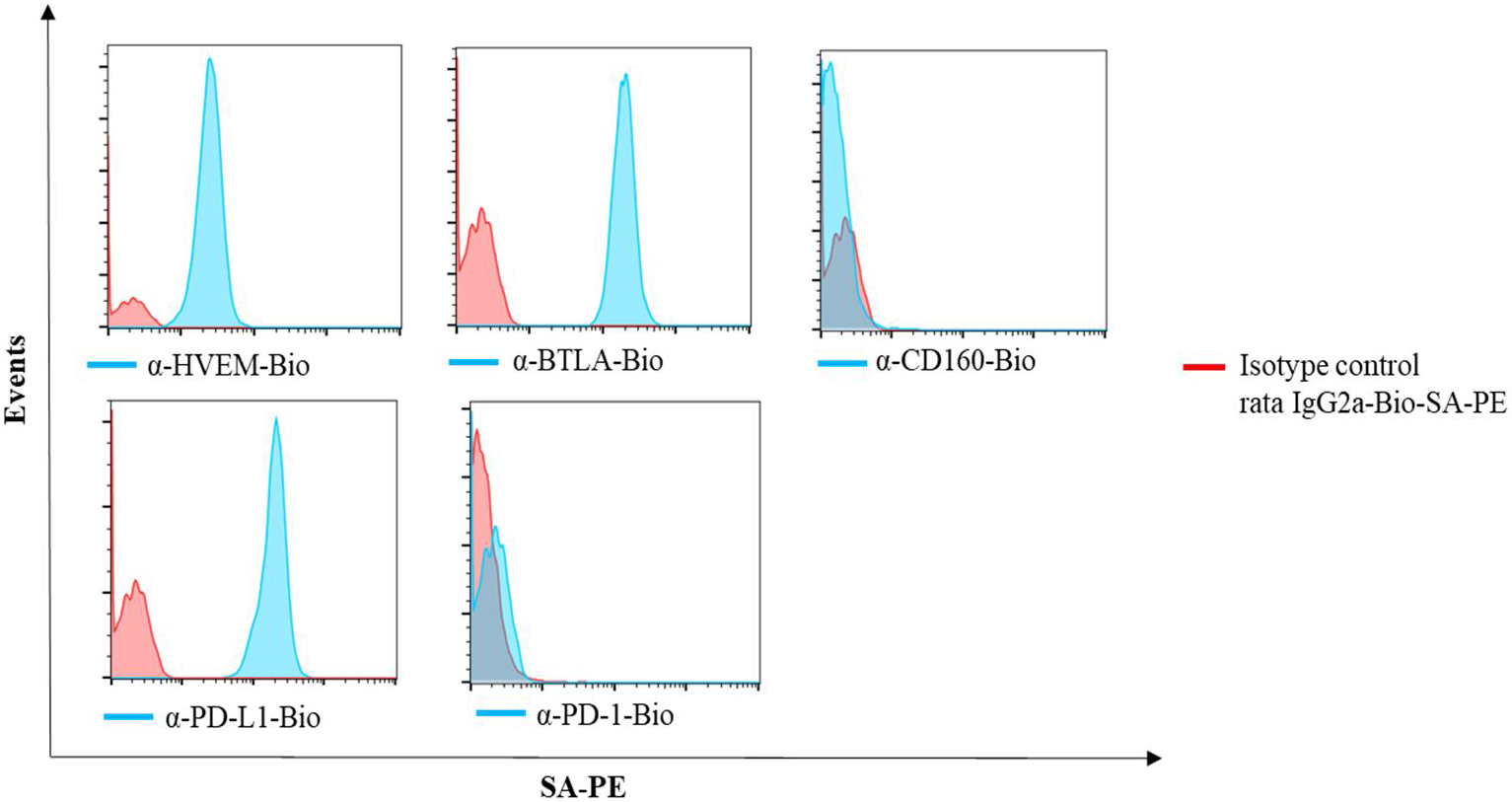
HVEM/BTLA/CD160 and PD-L1/PD-1 expression in A20 B-cell lymphoma line. In order to study HVEM, BTLA, CD160, PD-L1 and PD-1 expression by flow cytometry, biotinylated (Bio) monoclonal antibodies (mAbs) and SA-PE (blue) were used. We used biotinylated mouse anti-rat IgG2a as isotype control and SA-PE (red). The A20 B-cell lymphoma cells exhibited low expression of HVEM (MFI 25.7) and high expression of BTLA (MFI 177) and did not express CD160. On the other hand, A20 B-cell lymphoma cells expressed high levels of PD-L1 (MFI 178) but did not present PD-1 receptor expression. MFI: mean fluorescence intensity; IgG: immunoglobulin G, SA-PE: streptavidin-R-phycoerythrin conjugate; Bio: biotinylated.

### A20 tumour cells do not express mature B cell markers and show altered normal pattern expression of adhesion molecules and chemokine receptors

Our results demonstrated that A20 tumour cells expressed the typical B cell marker, CD19 (early B cell antigen), but they did not express markers of mature B lymphocytes such as CD21/35, IgM and IgD (Figure 3A). At the same time, the A20 B-cell lymphoma line showed an absence of molecules that play a critical role in B lymphocytes traffic from blood vessels to peripheral lymphoid organs. Such as chemokine (C-C motif) receptor 7 (CCR7) and adhesion molecules: LFA-1 (α_L_β_2_), because they lacked the β_2_ subunit (CD18), although they presented the α_L_ subunit (CD11a); and VLA-4 (α_4_β_1_) integrin, due to the absence of its α_4_ chain (CD49d) (Figure 3B).

**Figure 3.**
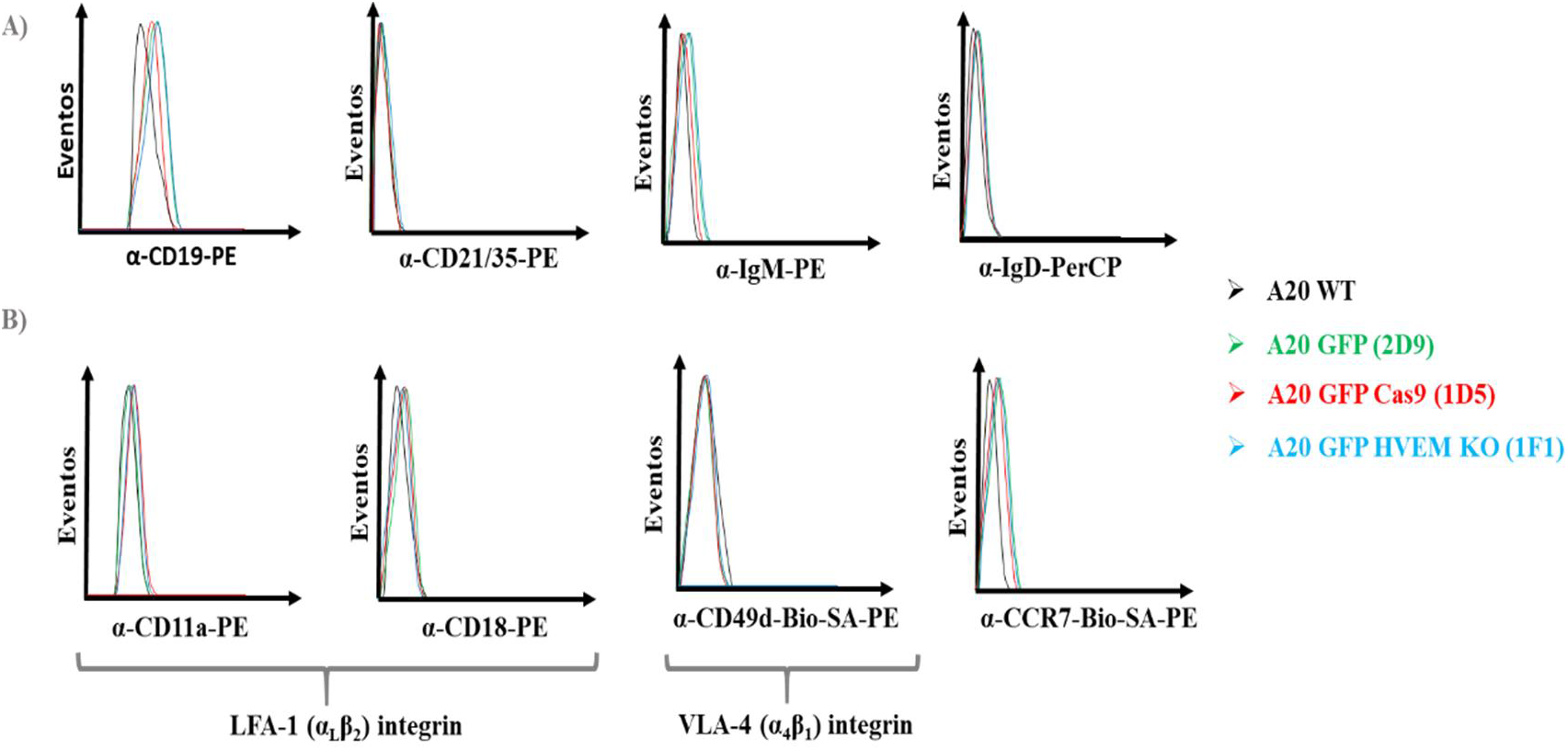
A20 tumour cells exhibit altered expression of mature B-cell markers, adhesion proteins and chemokine receptors. Representative histograms expression in different A20 B-cell lymphoma lines (A20 WT (black) as a control, A20 GFP (clone 2D9) (green), A20 GFP Cas9 (clone 1D5) (red), A20 GFP HVEM knock-out (KO) (blue)). (A) Study of CD19 (phycoerythrin (PE)), CD21/35 (PE), IgM (PE) and IgD (peridinin chlorophyll protein (PerCP)) expression in different A20 tumour cells. (B) Analyses of CD11a (α_L_ subunit) (PE) and CD18 (β_2_ subunit) (PE) expression (which form the LFA-1 (α_L_β_2_) integrin), and CD49d (α_4_ subunit) expression (which form part of VLA-4 (α_4_β_1_) integrin) in different A20 tumour cell lines.

### Genome editing of the first coding exon of HVEM by CRISPR-Cas9 exhibits two deletion forms, which leads to the loss of HVEM signal peptide expression

With the aim of distinguishing the host B-cell from A20 B-cell lymphoma, which was inoculated in Balb/c mice in order to cancer development and progression studies. We generated A20 cell line with stable eGFP marking (A20 GFP, clone 2D9) (Supplementary Figure S1). A20 GFP cells were used to obtain the A20 B-cell lymphoma HVEM knock-out (KO) line by CRISPR-Cas9 system.

As the target site of Cas9 nuclease, we established the first exon of HVEM sequence, which encodes together with part of the second exon for the HVEM signal peptide (1-38 aa) (Figure 4A). It was predicted with the help of *UniProt Signal Peptide* software. The main purpose was to determine whether the loss of HVEM signal peptide function prevented its membrane cell expression.

**Figure 4.**
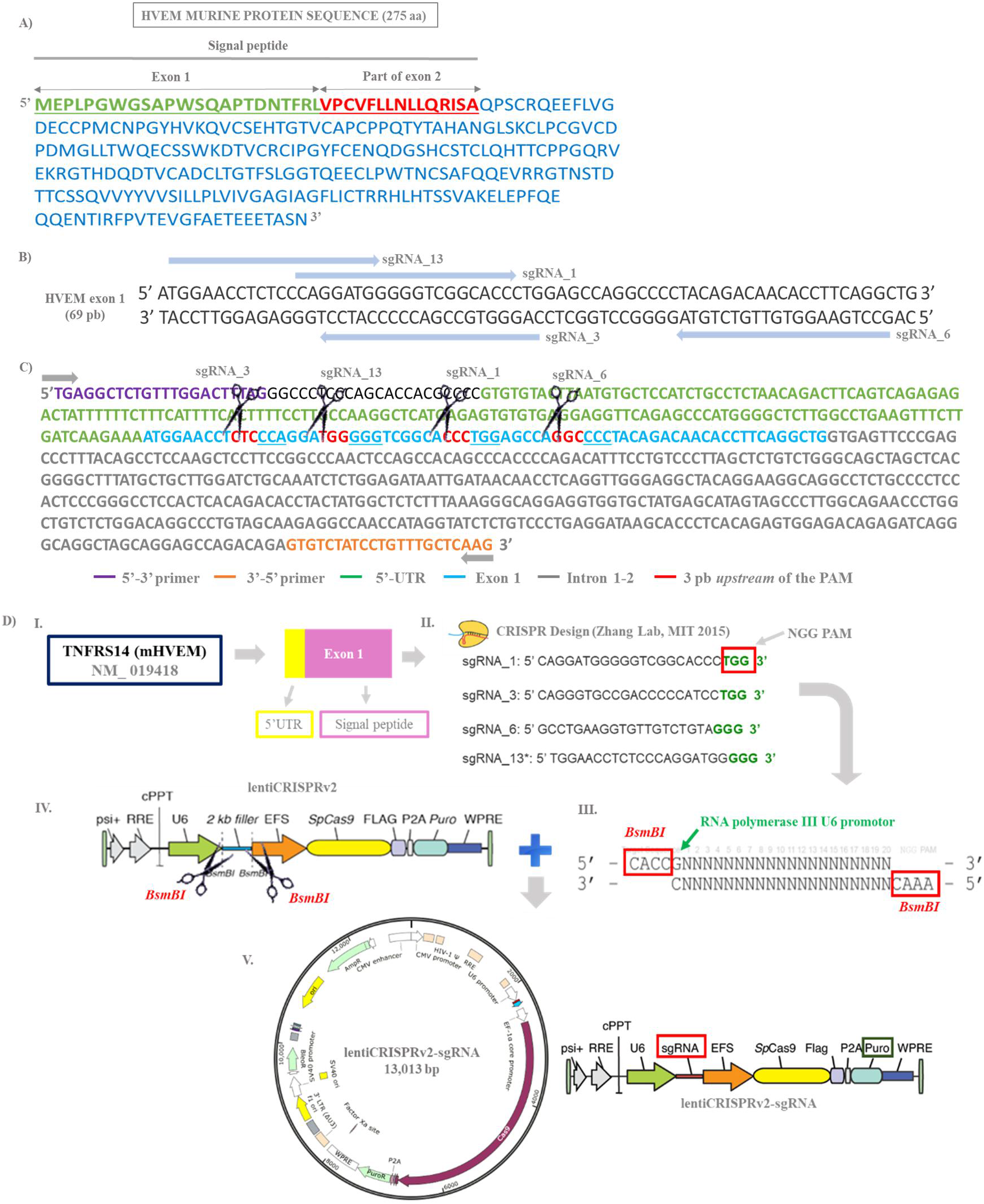
Design and selection of different sgRNAs that include the complete exon 1 of HVEM. Generation of lentiCRISPRv2-sgRNA (1, 3, 6 and 13) expression vectors. (A) Signal peptide delimitation in HVEM murine protein sequence (275 amino acids (aa)) (Balb/c strain). Signal peptide encompasses aa 1-38, and it is encoding by exon 1 (aa 1-23) (green and underlined) and part of exon 2 (aa 24-38) (red and underlined). The rest of the HVEM murine protein sequence is shown in blue. (B) Disposition of the selected sgRNAs (1, 3, 6 and 13) (blue arrows) on exon 1 of HVEM. (C) Cleavage sites of different sgRNAs selected on exon 1 (blue) of HVEM. Scissors mark the cleavage target site of Cas9 nuclease guided by each of the selected sgRNAs. Cleavages will occur 3 base pairs (bps) upstream (red) of the PAM sequence (underlined). (D) Cloning strategy for the construction of lentiCRISPRv2-sgRNA (1, 3, 6 and 13) expression vectors. I. A target site for the CRISPR-Cas9 system: exon 1 of HVEM (TNFRS14) sequence. II. sgRNAs designed and selected, having into account the presence of PAM sequence (NGG) (green). III. Generation double-stranded oligonucleotide (dsODN) duplex with matching overhangs (CACC/CAAA) (red box) that were resulting of *BsmBI* enzyme cleavage, and by adding guanine (G) necessary for RNA polymerase as transcription start. IV. Digestion of lentiCRISPRv2 vector with *BsmBI* enzyme. V. Individually ligation of each sgRNAs selected (1, 3, 6 and 13) into digested and purified lentiCRISPRv2 vectors, generating recombinant constructions that encode sgRNA and Cas9 nuclease (*sp*Cas9), and present a puromycin resistant cassette (PURO).

The different single-guide RNAs (sgRNAs) were designed from exon 1 of HVEM with *CRISPR Design* software, *Zhang Lab 2015*. We chose four of them (sgRNAs 1, 3, 6 and 13) (Figure 4B), in order to decide the best one to make a desirable mutation. These sgRNAs covered the whole exon 1 and cut at different positions inside of exon 1 (3 bp upstream of PAM) (Figures. 4B and 4C). The different sgRNAs (1, 3, 6 and 13) were individually cloned into a lentiCRISPRv2-sgRNA vector for co-expression with Cas9 nuclease (Figure 4D).

To select the best sgRNA for deletion of HVEM expression on the A20 cell surface, we used two different methods: T7/EI assay and pRR-eGFP system. In the first method, we transient transfected every single lentiCRISPRv2-sgRNA expressing sgRNA 1, 3, 6 or 13 and Cas9 nuclease in mammalian cells that expressed HVEM (Hepa 1-6), and examined the cleavage efficiency using T7/EI assay. The semi-quantification of T7/EI digestions revealed gene modification frequencies very similar for the different sgRNA studied (Supplementary Figure S2) (Supplementary Table S2). For the second technique, we used three different expression vectors: (i) pRR-eGFP vector, where was cloned the HVEM extracellular domain (ECD) (Supplementary Figure S3A), (ii) in order to separate the sgRNA and Cas9 expression, each sgRNA (1, 3, 6 and 13) was cloned into a sgRNA-tagRFP567 lentiviral vector coding for an APC fluorescence protein (Supplementary Figure S3B), and (iii) spCas9-PURO expressing Cas9 nuclease (Supplementary Figure S3C). This approach showed GFP positive HEK293T population percentages very similar for each sgRNAs examined (Supplementary Figure S4B). Therefore, we selected the sgRNA_1 to minimize undesired off-target cleavage, because it presented the highest on-target/off-target ratio (Table 1).

**Table 1.** Selection of sgRNA_1 presenting the lowest off-targeting activity. We obtained similar sgRNAs cleavage efficiencies with T7/EI assay and pRR-eGFP system. Both methods showed percentages of HVEM gene modification very similar to the different sgRNAs. We chose sgRNA_1 because presented the highest on-target/off-target marker (85%). This value is based on the difference of on-target activity, Cas9-mediated cleavage of target site guided by sgRNA, less off-target activities summation for each sgRNA. What it can resume in that formula*: off-targeting probability (%)* = *100%* − (∑ *% off*−*target activity for each sgRNA)*. Therefore, sgRNA_1 had the lowest probability of establishing complementarity in a non-specific area in the genome (red arrows).

| sgRNA | T7/EI assay | pRR-eGFP system | on-target/off-target ratio |
| --- | --- | --- | --- |
| ➡ sgRNA_1 | 26,80% | 25,97% | ➡ 85% |
| sgRNA_3 | 26,66% | 29,58% | 81% |
| sgRNA_6 | 26,44% | 28,14% | 69% |
| sgRNA_13 | 25,20% | 24,82% | 55% |

In order to get HVEM deficient A20 GFP cell line, these cells were electroporated with the lentiCRISPRv2-sgRNA_1 (coding for both sgRNA_1 selected and Cas9 nuclease expression). PCR amplification and sequencing analyses of HVEM target site from heterogeneous electroporated A20 GFP cells with CRISPR-Cas9 system and sgRNA_1 selected, allowed us to detect two different deletions mutated forms of HVEM (100 and 300 bp HVEM deletion forms) (Figure 5A) (Supplementary Figures S5A and S5B). Both HVEM deletions were characterized by the disappearance of complete exon 1 (target of the CRISPR-Cas9 system). The truncated mutation of HVEM of 100 bp was characterized by presenting part of the 5’UTR sequence (35 bp of the total 156 bp) and part of the intron 1-2 sequence (66 bp of the total 731 bp), and for the complete loss of exon 1 (Supplementary Figure S5A). The truncated mutation of HVEM of 300 bp presented 242 bp of the total 731 bp of intron 1-2, having completely lost 5’UTR and exon 1 (Supplementary Figures S5B). The clones (1F1 and 4D10) selected by limiting dilution from A20 GFP electroporated heterogeneous population exhibited the 100 bp deletion form and did not present Cas9 nuclease genomic integration (Figures 5B and 5C).

**Figure 5.**
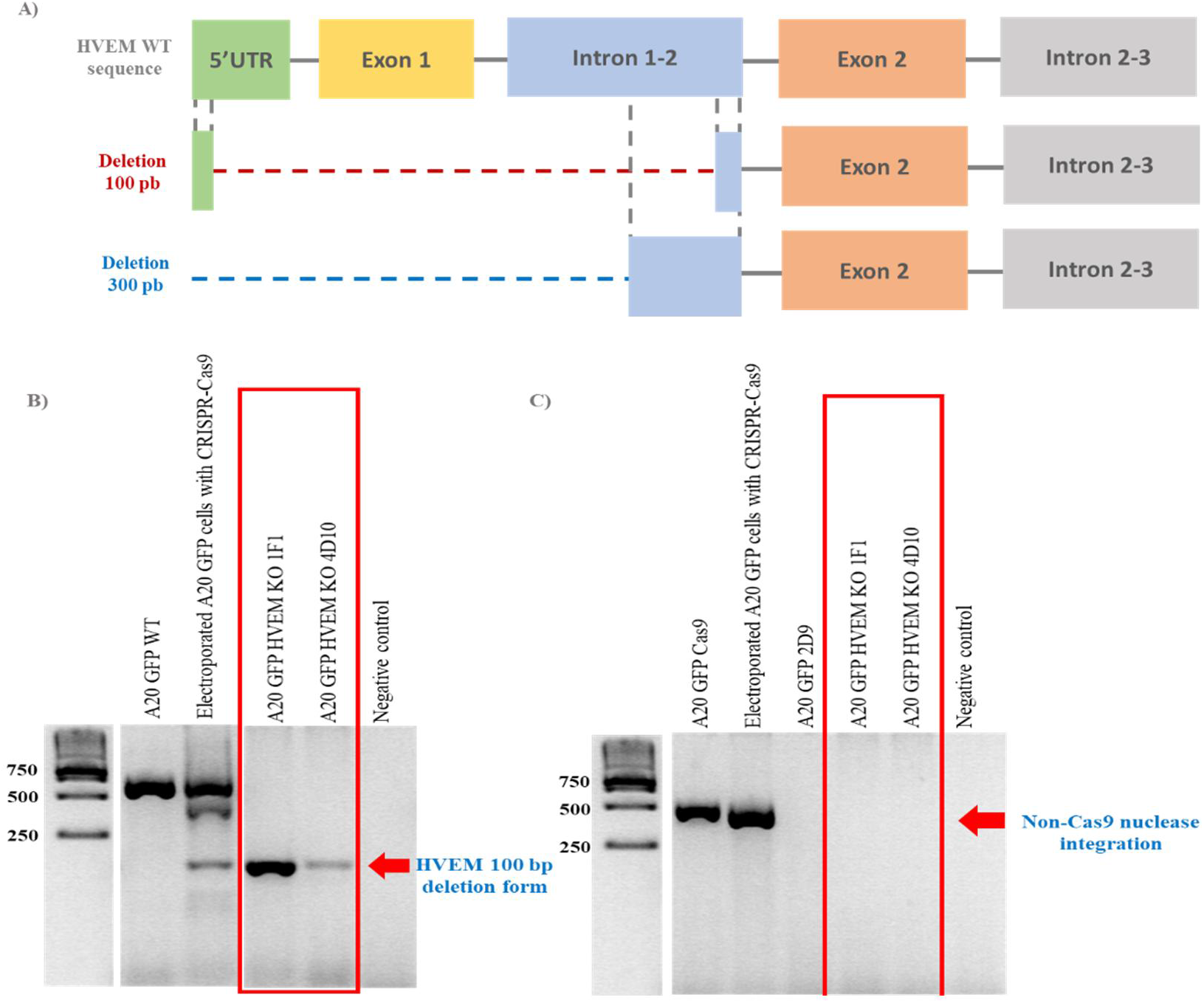
Schematic comparison between HVEM WT sequence and the resultant HVEM deletion forms by CRISPR-Cas9-sgRNA_1 editing. PCR amplification results of HVEM target region of the editing system and Cas9 nuclease integration. (A) Dotted lines (red for 100 bp deletion and blue for 300 bp deletion) indicate the lost regions of each HVEM truncated forms Cas9 nuclease-mediated guided by sgRNA_1. (B) Amplification by PCR of HVEM target site of Cas9 cleavage and sgRNA_1. Heterogeneous A20 GFP electroporated with CRISPR-Cas9-sgRNA_1 population showed both HVEM deletions forms (100 and 300 bp) and HVEM WT forms. However, the two clones selected (A20 GFP HVEM knock-out (KO), clones 1F1 and 4D10) from this heterogeneous population of electroporated A20 GFP cells exhibited the truncated form of 100 bp HVEM. A20 GFP WT cells were used as a positive control of HVEM WT amplification. (C) Cas9 nuclease amplification by PCR allowed us to know that A20 GFP HVEM KO clones (1F1 and 4D10) not presented Cas9 nuclease integration, despite the fact that the heterogeneous population of cells from which it started had the Cas9 nuclease. The reason could be that Cas9 integration phenomenon would have occurred in a very small percentage, which we could detect by PCR. As a positive control of Cas9 amplification, we used the Cas9 stable expression A20 GFP line (clone 1D5) generated by lentiviral systems.

### The loss of HVEM signal peptide expression by CRISPR-Cas9-sgRNA_1 system is sufficient to prevent membrane HVEM expression on A20 GFP cells

The complete loss of HVEM exon 1 by CRISPR-Cas9-sgRNA_1 in both types of deletion forms (100 and 300 bp) resulted in the change of open reading frame (ORF), which began in methionine (M) residue situated at position 56. This led to the loss of HVEM signal peptide expression, and also the loss of a part of the coding extracellular domain (Figure 6A). In order to prove whether the loss of HVEM signal peptide expression prevented HVEM surface expression on A20 GFP clones 1F1 and 4D10 selected (which contained 100 bp HVEM deletion form), we analysed the HVEM expression by flow cytometry with biotinylated mouse anti-HVEM monoclonal antibody (mAb) (clone 6C9) and SA-PE. Our data revealed that A20 GFP clones 1F1 and 4D10 showed an HVEM expression pattern very similar to anti-rat IgG_2a_ isotype control (Figure 6B). In contrast to the A20 GFP WT cell line (clone 2D9), which was positive for HVEM expression (Figure 6B). These data confirmed that the absence of HVEM signal peptide expression was sufficient to complete abrogation of membrane HVEM expression on A20 GFP tumour cells. In addition, it suggested that mutations were bi-allelic. Therefore, we generated an HVEM deficient A20 B-cell lymphoma line (A20 GFP HVEM KO), and select the clone 1F1 because it presented the best in vitro growth condition.

**Figure 6.**
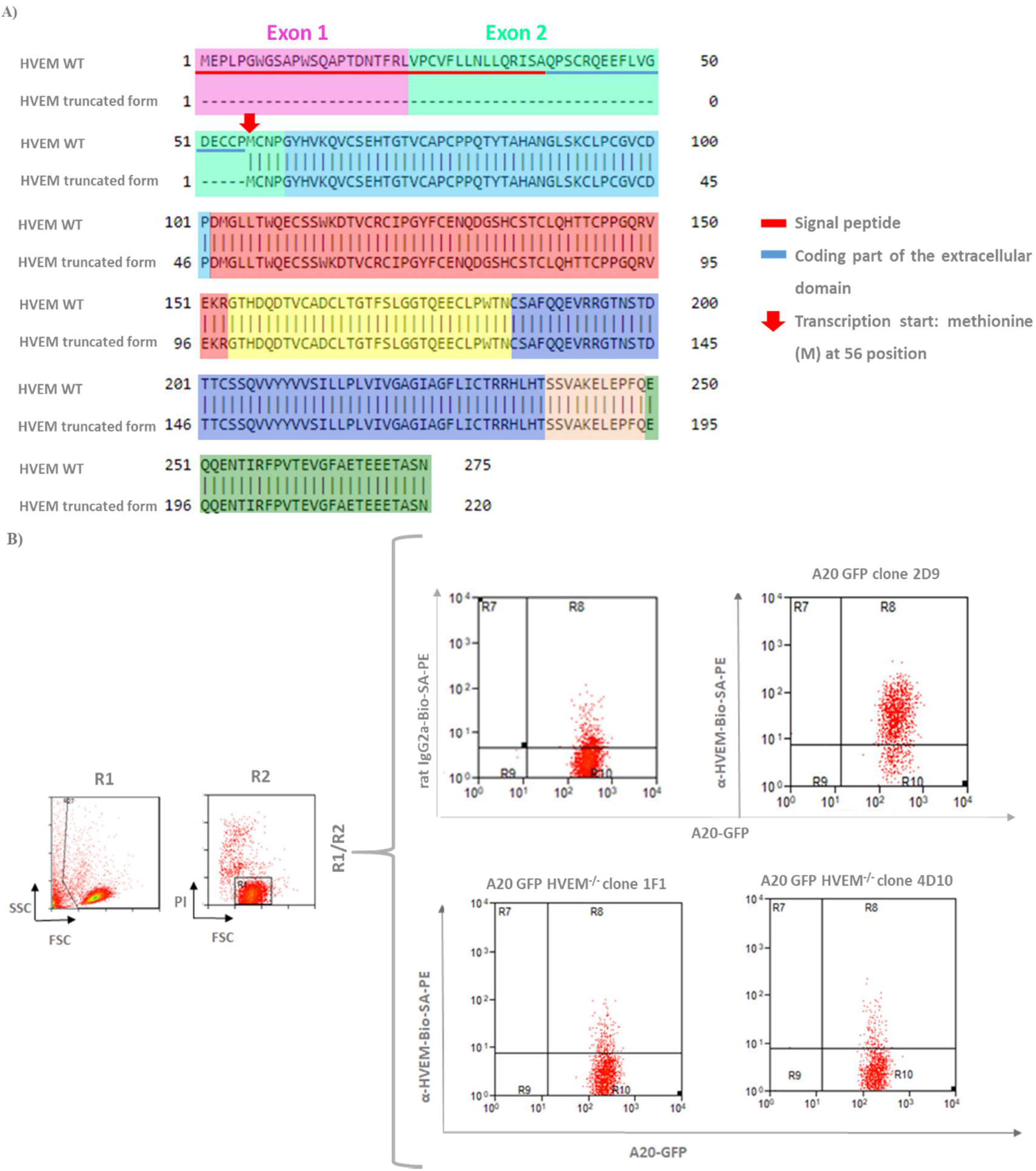
Complete deletion of HVEM exon 1 by CRISPR-Cas9-sgRNA_1 system results in the loss of signal peptide expression, which leads to complete abrogation of membrane HVEM expression in A20 GFP tumour cells. (A) The protein sequence comparison between HVEM WT (top) and HVEM deletion form (bottom) determined that truncated HVEM protein by CRISPR-Cas9-sgRNA_1 had lost signal peptide expression (underlined in red) (which is coding by exon 1 (pink) and part of exon 2 (light green), because of the absence of exon 1 changed the initial methionine (M) residue to position 56 (red arrow). This change in the open ready frame (ORF) prevented the signal peptide expression and the expression of an extracellular domain part (underlined in blue). The different colours mean the 8 exons that makeup an HVEM protein sequence. (B) We confirmed by flow cytometry that the loss of signal peptide expression was sufficient to prevent HVEM expression on A20 GFP surface (A20 GFP HVEM^−/−^ clones 1F1 and 4D10). In order to check HVEM expression, we used biotinylated monoclonal anti-HVEM and SA-PE. As isotype control, we used biotinylated mouse anti-rat IgG2a and SA-PE. As a positive control of HVEM expression were used A20 GFP cells (clone 2D9). Abbreviations R1: initial population, R2: living cell population. FSC: forward scatter, SSC: side-scatter and PI: propidium iodide.

### The importance of non-integration of Cas9 nuclease: CD40 expression increase in Cas9 transduced A20 GFP cells

To evaluate the possible consequences of stable Cas9 expression in A20 tumour cells, we generated an A20 GFP cell line with constitutive Cas9 expression by lentiviral vectors (A20 GFP Cas9, clone 1D5). We demonstrated by flow cytometer with anti-CD40 mAb that A20 GFP Cas9 cells showed higher CD40 (co-stimulatory receptor) expression in compare with A20 cell lines (A20 WT, A20 GFP (clone 2D9), and A20 GFP HVEM KO (clone 1F1)), which did not present Cas9 genomic integration (Figure 7A). This result suggests the Cas9 nuclease pathogen-associated molecular patterns (PAMPs) recognition by A20 tumour cells (Hua & Hou 2013) (Jiang et al. 2007) (Figure 7B). This data highlights the importance of non-integration of Cas9 in the A20 cell genome to avoid secondary undesirable effects, such as a high in vivo immune response against the tumour.

**Figure 7.**
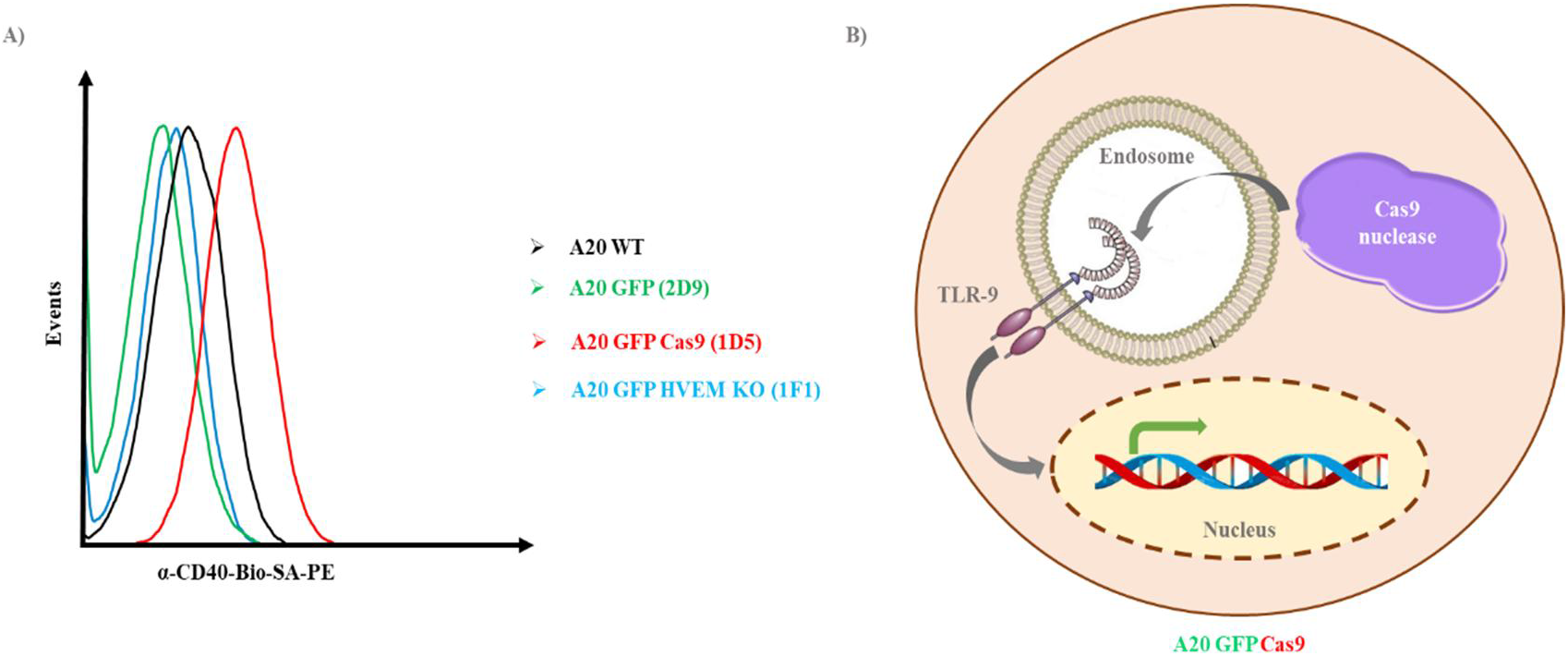
Modulation of CD40 co-stimulator receptor expression in the A20 tumour line in response to transduction with Cas9 nuclease. (A) A20 GFP with Cas9 nuclease stable expression (A20 GFP Cas9, clone 1D5) (red) showed a higher expression levels of CD40 co-stimulator receptor, in comparison with the others cell lines that do not express Cas9 (A20 WT (black), A20 GFP, clone 2D9 (green), A20 GFP HVEM KO, clone 1F1 (blue)). Biotinylated monoclonal anti-CD40 and SA-PE were used to study CD40 expression levels. (B) Schematic drawing of Cas9 nuclease pathogen-associated molecular patterns (PAMPs) recognition by Toll-like receptor 9 (TLR-9), which would lead to activation of gene expression like the one that codes for CD40 receptor.

### The RNA-guided 1 (sgRNA_1) selected does not present off-target activity detectable

Off-target cleavage is based on the proximity of non-complementary nucleotides to PAM; that is, Cas9 nuclease will cleave worse non-complementary nucleotides situated closer to 3’ N-terminal (Figure 8A). To determine whether the sgRNA_1 selected showed off-target activity, we amplified by PCR and sequenced the first five sequences of genes with more off-target probability from A20 GFP HVEM^−/−^ (1F1) genome (Supplementary Table S1) (Figure 8B): *Hpgd* gene (NM_008278), *Gpsm1* gene (NM_153410), *Sipa1l3* gene (NM_001081028), *Neto1* gene (NM_144946) and *Oplah* gene (NM_153122). *Hpgd*, *Gpsm1* and *Neto1* genes are intronic sequences, while *Sipa1l3* and *Oplah* genes are exonic sequences. Then, sequenced off-target genes were compared with their respective WT sequences revealing that they had not been modified by CRISPR-Cas9-sgRNA_1 off-target cleavage (Figure 8C).

**Figure 8.**
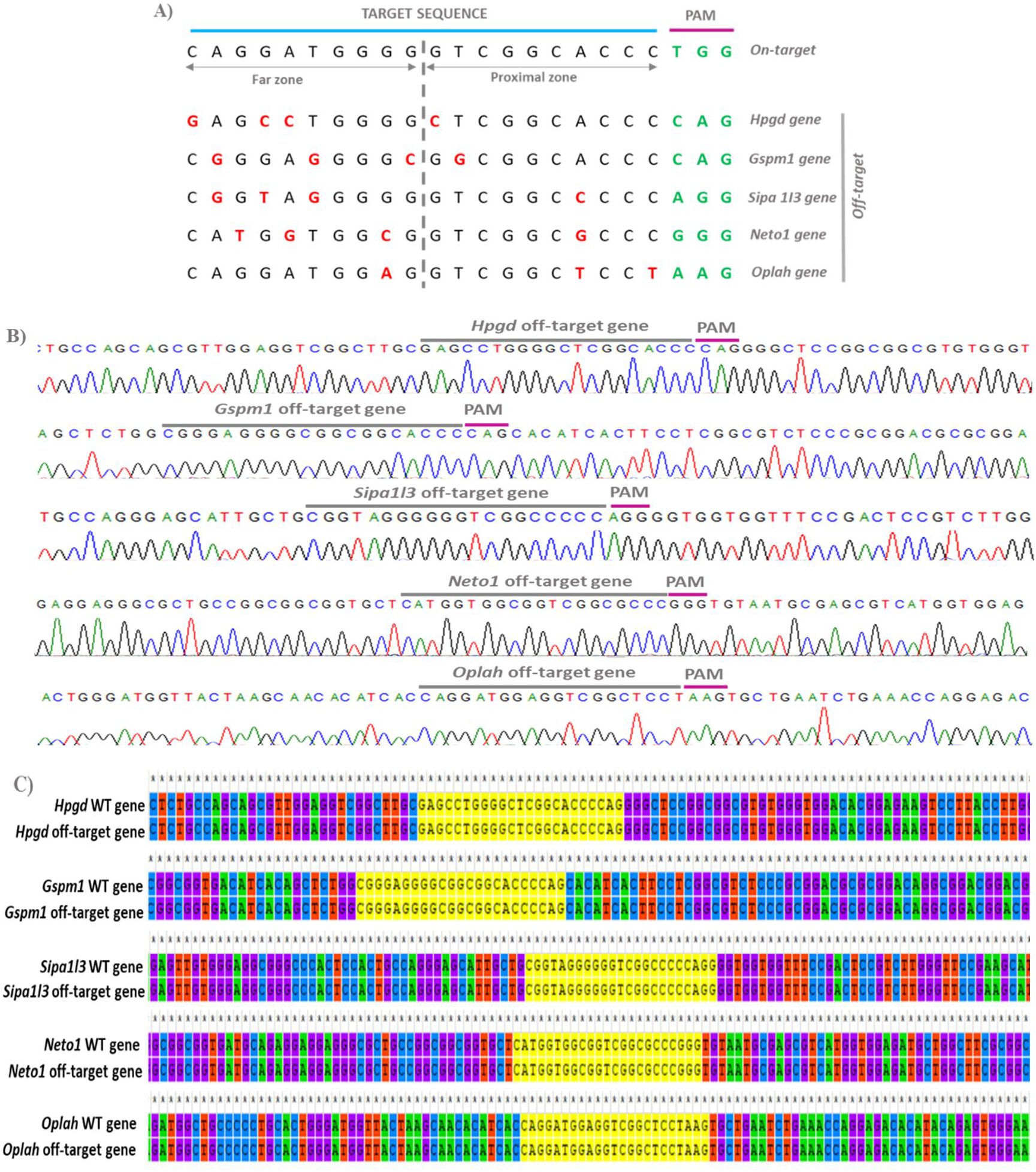
Analysis of the potential sgRNA_1 off-target activity in A20 GFP HVEM^−/−^ (1F1) genome. (A) Position of the different non-complementary nucleotides in each off-target sequences. The off-target of probability depends on mismatches position (red) to the PAM sequence (green). The non-complementary nucleotides that are in the proximal zone (8-10 bp upstream to PAM) reduce the possible off-target activity of Cas9 nuclease guided by sgRNA_1. (B) Result of off-target genes sequencing from A20 GFP HVEM^−/−^ (1F1) genome. (C) Comparison between the different off-target genes sequenced and its corresponding WT sequences. In yellow is indicated that the different off-target sequences had not been modified by CRISPR-Cas9-sgRNA_1.

### The loss of HVEM expression does not lead to modification of A20 tumour cells in vitro duplicate rate but provokes a decline of BTLA expression

With the objective to determine whether the absence of membrane HVEM expression in A20 tumour cells affected in vitro replication capacity, we realized A20 GFP WT and A20 GFP HVEM^−/−^ cell counting at 24 and 48 hours in culture and exponential growth. We found that there were not any significant differences between A20 GFP WT and A20 GFP HVEM^−/−^ duplicate rates (Supplementary Table S3). This result suggests that the loss of HVEM expression does not affect to A20 tumour cell in vitro proliferative ratio. At the same time, A20 GFP HVEM^−/−^ exhibited a lower BTLA expression in compare with A20 GFP WT (Figure 9A), what may be due to the impossibility to form of stable *cis* HVEM-BTLA complex on the A20 GFP HVEM^−/−^ surface (Figure 9B).

**Figure 9.**
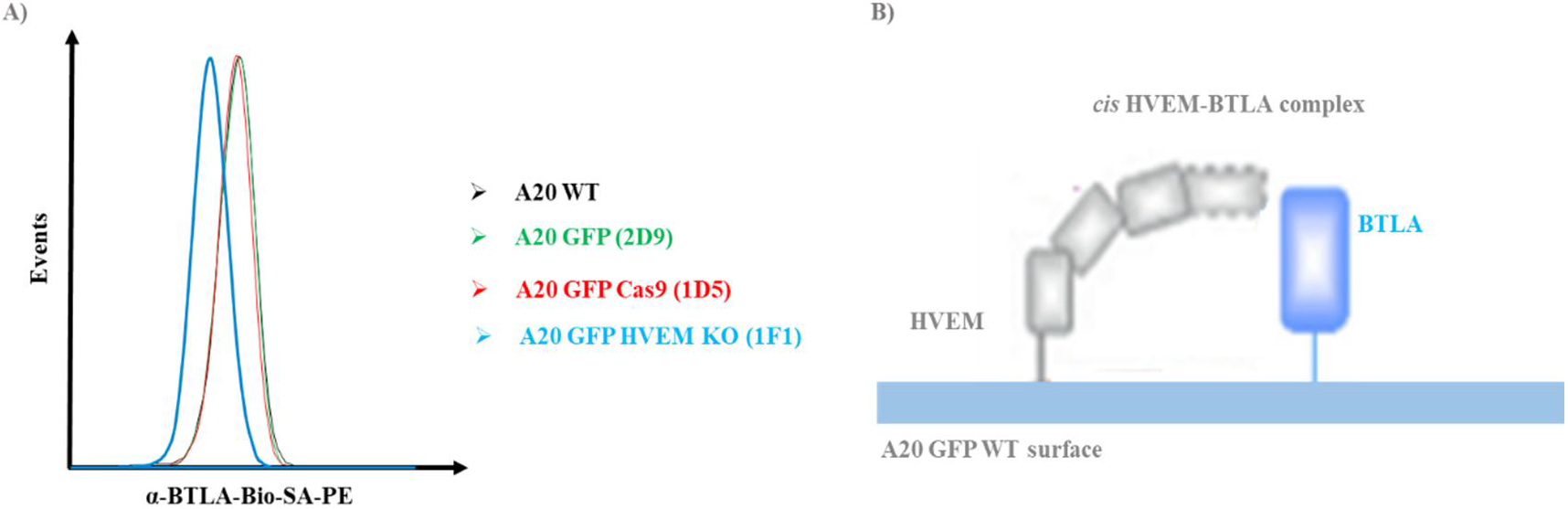
The loss of HVEM expression in the membrane of A20 GFP HVEM^−/−^ cells reduces BTLA receptor expression levels. (A) A20 GFP HVEM knock-out (KO) (clone 1F1 (blue)) showed lower BTLA expression level than the others analysed cell lines (A20 WT (black), A20 GFP (clone 2D9) (green) and A20 GFP Cas9 (clone 1D5) (red)). Biotinylated monoclonal anti-BTLA and SA-PE were used. (B) Graphical representation of *cis* HVEM-BTLA interaction on A20 GFP WT surface cell.

### The absence of HVEM expression in A20 tumour cells does not provoke changes in metastatic dissemination pattern to different lymphoid organs

In order to study whether the loss of HVEM expression on A20 lymphoma cells membrane provoked changes in its dissemination capacity to primary and secondary lymphoid organs, we analysed by flow cytometer the proportion of A20 GFP WT and A20 GFP HVEM^−/−^ in these hematopoietic organs at days 22, 28 and 34 post-tumour implantations. The A20 lymphoma colonization of lymphoid organs was only possible from the intermediate phase of tumour development (day 28 post-implantation of tumour) in case of the spleen, lymph nodes and bone marrow (Figure 10). We did not saw any significant differences between A20 GFP WT and A20 GFP HVEM^−/−^ metastatic dissemination to different lymphoid organs (Figure 10). These data suggest that the loss of HVEM expression does not alter the colonization efficiency of A20 tumour cells to different hematopoietic compartments.

**Figure 10.**
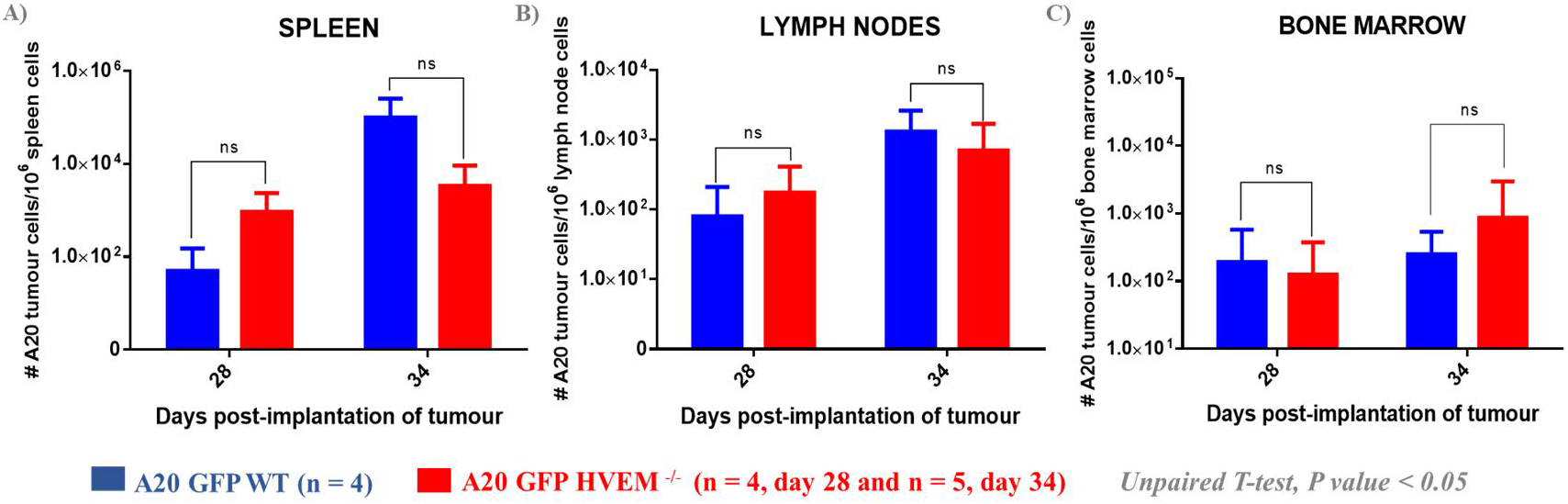
The efficiency of the spleen, lymph nodes and bone marrow colonization by A20 GFP HVEM^−/−^ compare with A20 GFP WT tumour cells. Representation of A20 tumour cells (blue: A20 GFP WT (n = 4 at both days 28 and 34 post-implantation of tumour) and red: A20 GFP HVEM^−/−^ (n = 4 at day 28 and n = 5 at day 34 post-implantation of tumour) absolute number (#) per one million of the corresponding lymphoid organ (spleen (A), lymph nodes (B) and bone marrow (C)). The results are expressed as mean±SD (standard deviation). ns: no significant differences (unpaired T-test, p-value < 0.05).

### Thymus colonization by A20 lymphoma cells in advanced stages of cancer progression promotes an endogenous B cells expansion

A20 lymphoma cells were only capable of thymus colonization in advantage stages of cancer development (at day 34 post-implantation of tumour) (Figure 11A), unlike what occurred in the rest of lymphoid organs. Moreover, we observed other curious data in this hematopoietic compartment. In homeostatic conditions, the normal percentage of thymic B cells are less than 0.5%. However surprisingly, the number of thymic B cells began to increase in mice inoculated with both types of lymphoma cells, A20 GFP WT and A20 GFP HVEM^−/−^ (Figure 11B). Even before thymus colonization by A20 lymphoma cells was evident (at day 28 post-implantation of tumour), the proportion of endogenous B cells showed an unusual increment (Figure 11B). However, it was in the final stages of the illness when the number of endogenous B cells reached a higher percentage (Figure 11B). This data suggests that changes would be taking place in the thymus microenvironment, which would promote endogenous B cells expansion and thymus implantation by A20 lymphoma cells.

**Figure 11.**
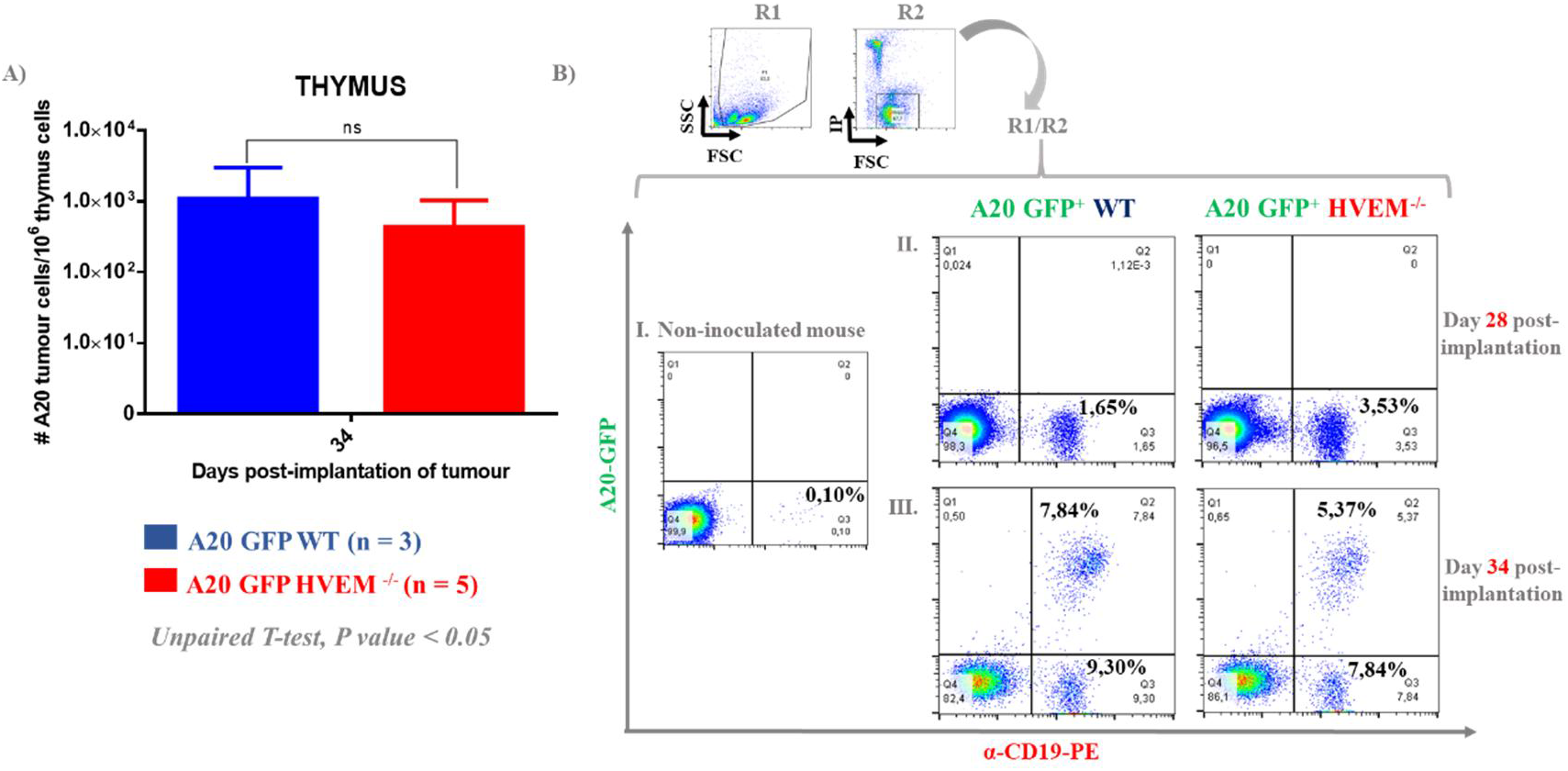
Thymus colonization kinetics by A20 GFP HVEM^−/−^ and A20 GFP WT. The peculiar increase of endogenous B cell in the thymus of mice inoculated with both A20 tumour cells. (A) Representation of A20 tumour cells absolute number (#) per 1 × 106 thymic. The results are expressed as mean±SD. No significant differences were observed (ns) (unpaired T-test, p-value < 0.05) between A20 GFP WT (blue, n = 3) and A20 GFP HVEM^−/−^ cells (red, n = 5). (B) I. In the thymus of a control mouse (not inoculated with the A20 tumour line), the population of B cells (CD19^+^ GFP^−^) was ≈ 0.1%. II. At day 28 post-inoculation of A20 GFP WT and A20 GFP HVEM^−/−^ lines, in the thymus, there was an increase in the number of B cells with respect to B cell population of a control mouse thymus. III. At day 34 post-inoculation, when A20 GFP WT and A20 GFP HVEM^−/−^ cells were detected in the thymus, the highest percentages of endogenous B cells appeared. R1: initial population gate and R2: living cells gate.

### The loss of HVEM expression on A20 lymphoma cell alters the capacity of immune cells recruitment to hepatic tumour nodules

Hepatic tumour nodules created by A20 lymphoma cells were used to study the tumour ability to the recruitment of the immune cells, after having lost the membrane HVEM expression (Figure 12A). The main reason to use hepatic tumour nodules is that lymphoid organs did not permit us to distinguish between resident immune cells and infiltrating leukocytes in the tumour (TILs). At the intermediate phase of cancer progression (day 28), the TILs (CD8 and CD4 T cells, B cells, NK cells and CD11b myeloid cells) number in hepatic tumour nodules formed by A20 GFP HVEM^−/−^ was significant lower than TILs attracted to A20 GFP WT hepatic nodules (Figures 12B and 12C). Consequently, the A20 tumour cells proportion was higher in A20 GFP HVEM^−/−^ hepatic nodules (Figure 12B). These preliminary studies suggest that the loss of HVEM expression could be altering the efficiency of immune cell recruitment to the tumour microenvironment. However, these significant differences in the TILs recruitment ability by tumour were not observed at least stages of lymphoma (34 days) (data not showed).

**Figure 12.**
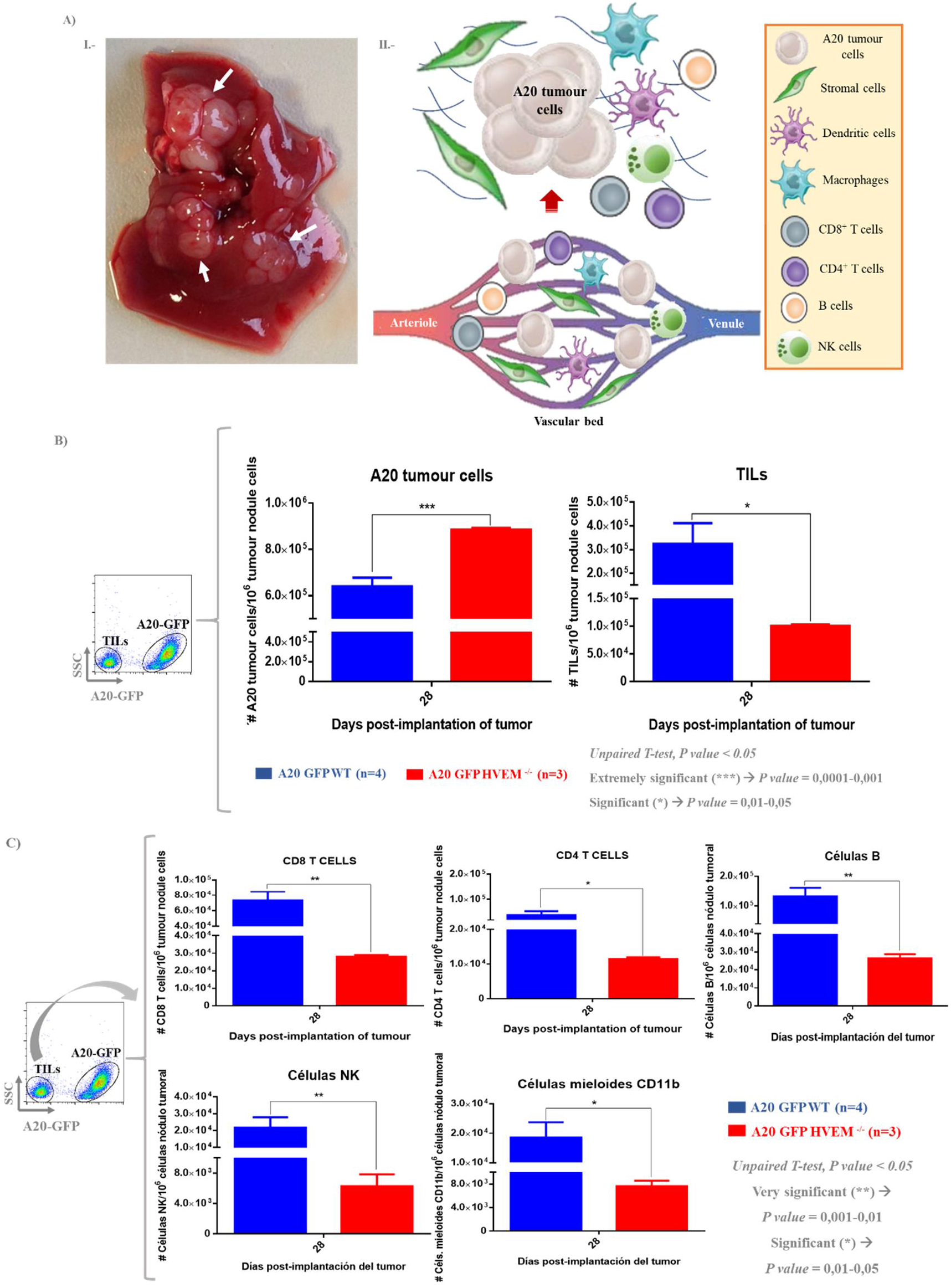
Study of the immune system cells recruitment by A20 GFP HVEM^−/−^ compare with A20 GFP WT in hepatic tumour nodules at day 28 post-tumour implantation. (A) I. White arrows indicate the hepatic tumour nodules formed by A20 tumour cells. II.-Graphical representation of a possible tumour microenvironment formed by immune system cells and non-hematopoietic cells, like stromal cells. (B) Representation of A20 tumour cells and infiltrating leukocytes in the tumour (TILs) absolute number (#)/ 1 × 106 hepatic tumour nodule cells at day 28 post-implantation. (C) Representation of different immune cells absolute number (#)/1 × 10^6^ hepatic tumour nodule cells. Blue: control group inoculated with A20 GFP WT line and red: problem group inoculated with line A20 GFP HVEM^−/−^. The statistical differences resulting from an unpaired Student’s t-test are indicated as: extremely significant (***) (p-value = 0.0001-0.001), very significant (**) (p-value = 0.001-0.01), and significant (*) (p-value = 0.01-0.05).The results are shown as mean±SD.

### A20 lymphoma cells exert an immune cells exclusion phenomenon

We analysed the A20 tumour cells (both A20 GFP WT and A20 GFP HVEM^−/−^) infiltration in hepatic tumour nodules by classical immunohistology in order to know their distribution. A20 GFP WT distribution could be diffuser; meanwhile, A20 GFP HVEM^−/−^ infiltration seemed to have a nodular pattern (Figures 13A, 13B, 13C and 13D). However, these differences were not clear. What is clear is that A20 tumour cells were able to exclude TILs. TILs appeared around of lymphoma, instead of intermingling with them (Figures 13E and 13F). It may be an immune evasion mechanism by A20 B-cell lymphoma, limiting the possible interactions between effector immune cells and A20 tumour cells that they led to their destruction.

**Figure 13.**
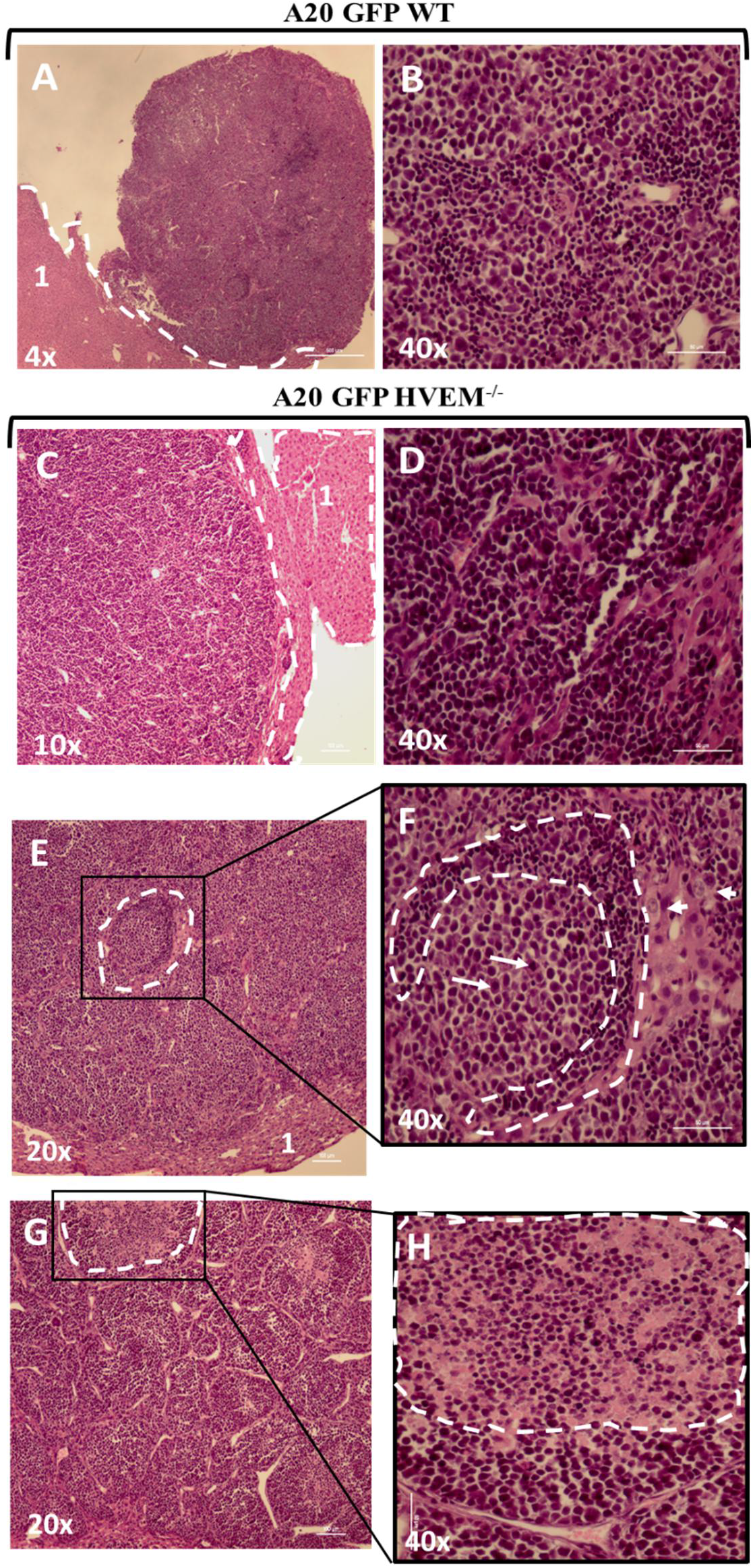
Distribution of A20 tumour cells and TILs in the hepatic tumour nodules. Hepatic tumour nodules formed by A20 GFP WT line (A) and A20 GFP HVEM^−/−^ line (B) at day 28 post-implantation of the tumour. Infiltration pattern of A20 GFP WT (C) and A20 GFP HVEM^−/−^ (D) in tumour nodules. Liver tissue is delimited by a white dotted line. (E) (F) Distribution of A20 tumour cells and infiltrating leukocytes in the tumour (TILs). (F) TILs (smaller cells than the large neoplastic A20 cells) are distributed around the tumour (area delimited by a white dotted). The short arrows indicate the A20 tumour cells with centroblastic morphology and the long arrows indicate the tumour cells with immunoblastic morphology. (G) Structural organization of the tumour nodules of the liver (necrotic dotted area in white (H)).

At the same time, we observed that A20 tumour (both A20 GFP WT and A20 GFP HVEM^−/−^) nodules formed in lymph nodes compressed secondary B-cell follicles in the cortex area of the lymph nodes, and they lost their spherical shape to acquire a more oval shape. Similar to occur in hepatic tumour nodules, TILs were disposed around A20 tumour nodules (Figure 14).

**Figure 14.**
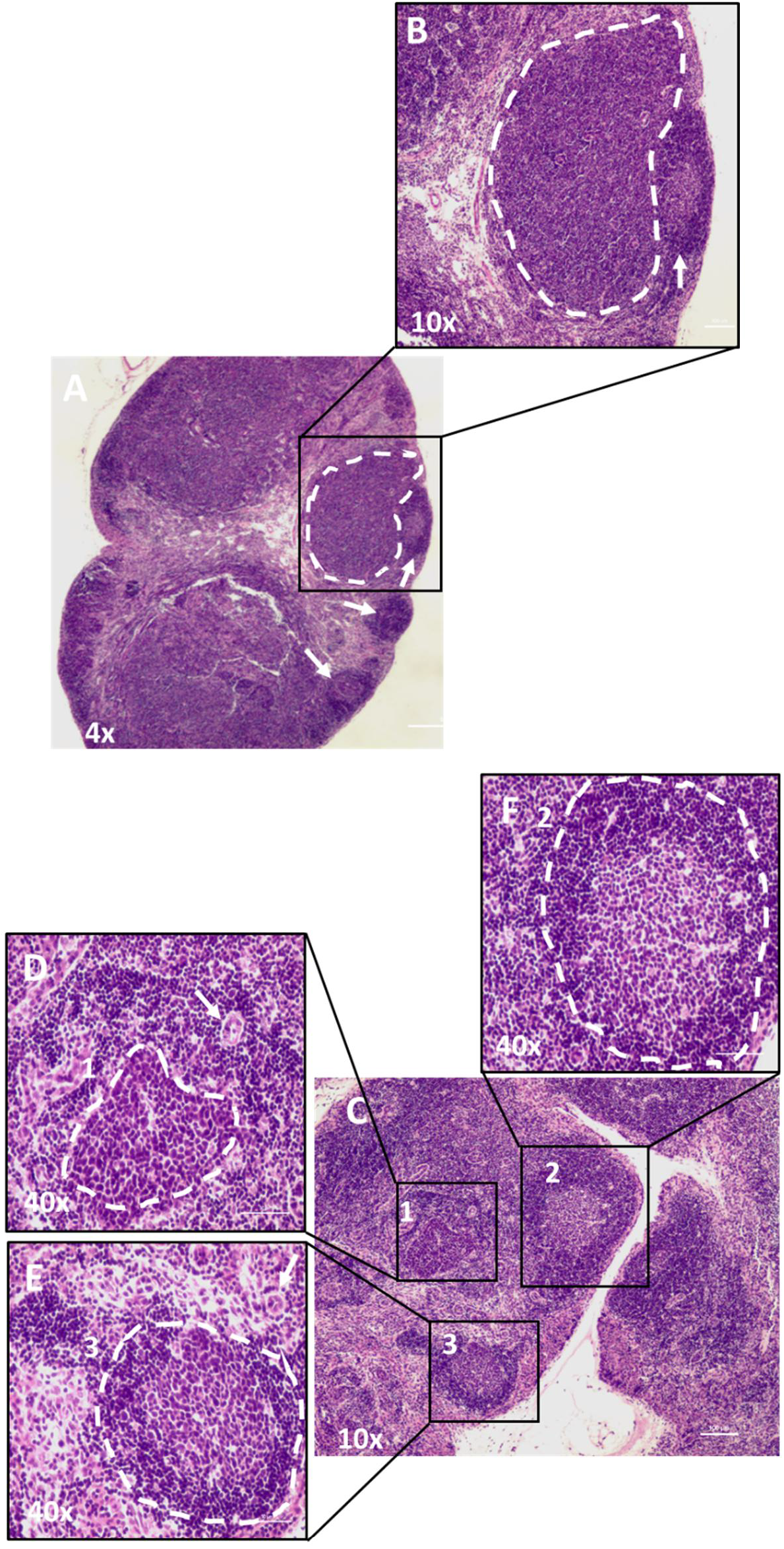
Infiltration of A20 tumour cells in the lymph nodes at day 28 post-implantation. (A) (B) A20 tumour cells (white dotted area) compress the follicles on the lymph node cortex acquiring a more oval shape (white arrow). (C) Image showing 1A20 tumour cells and 2,3follicles with normal morphology. (D) Infiltration of A20 tumour cells (dotted area in white) forming a mass and the TILs distributed around it. (E) (F) B cell follicles with normal morphology. The clearest part of the follicle is the germinal centre (GC), in which B cells proliferate and differentiate into plasma cells that produce antibodies. White arrows indicate the high endothelial venules (HEVs) (cross-section). The HEVs present unusually high endothelial cells and through them enter the lymphocytes from the blood to the lymph node.

## DISCUSSION

### Molecular and in vitro consequences of the loss of HVEM in A20 B-cell lymphoma line surface

The intrinsic result of the loss of HVEM expression on the A20 B-cell lymphoma surface was the decline in BTLA expression. The reason may be the impossibility to form the cis HVEM-BTLA complex, which would lead to down-regulation of BTLA expression. This interaction is very stable, and its dissociation only can be mediated by the membrane-bound form of LIGHT in trans, whose expression is induced after immune response activation (Pasero et al. 2012; Steinberg et al. 2011; Ware & Sedy 2011). The decrease in BTLA expression levels had also been observed in FL that presented *TNFRSF14* loss-of-function mutations; however, it had not been proved yet in DLBCL (Boice et al. 2016; Pasqualucci & Dalla-Favera 2018). These suggest that HVEM expression condition the BTLA expression in the same cell surface.

The signalling through HVEM receptor mediated by its BTLA ligand in trans induces the expression of survival and proliferation genes by activating of NF-κB transcription factor (McGrath & Najafian 2012; Adam et al. 2003; Murphy et al. 2006). For this reason, it could have been possible that A20 GFP HVEM^−/−^ exhibited a different in vitro duplication rate in compare with A20 GFP WT. However, this was not the case. This may be due to HVEM (expressed in A20 GFP WT) was mostly forming the cis HVEM-BTLA complex, preventing the interaction HVEM-BTLA in trans, limiting the NF-κB activation through HVEM pathway in A20 tumour cells (Shui et al. 2011; Steinberg et al. 2011; Ware & Sedy 2011). If so, the loss of HVEM expression in A20 tumour cells would disrupt the cis HVEM-BTLA interaction, which would limit the inhibitory effects in downstream of B cell antigen receptor (BCR) signalling. However, in vitro BCR stimulation would be absent, as well as, the possible co-stimulatory action of CD4 T-helper cells (which would take place in vivo). Therefore, one possible explanation could be that the main role of HVEM in these A20 tumour cells is to act as a ligand of its co-inhibitory BTLA receptor expressed in the same cell.

### The implication of the loss of HVEM in the metastatic A20 B-cell lymphoma spread to hematopoietic organs

The fact that there had not any significant differences in the efficacy of metastatic colonization to primary and secondary lymphoid organs by A20 GFP HVEM^−/−^ in compare with A20 GFP WT suggests that HVEM is not implicated in the A20 lymphoma cells dissemination, and other molecules govern A20 tumour cells homing to hematopoietic organs.

A20 tumour cells did not express CC-chemokine receptor 7 (CCR7), which is crucial in B-cell lymphoma dissemination to lymph nodes (Pals et al. 2007; Till et al. 2014; Scott & Gascoyne 2014). Neither LFA-1 (α_L_β_2_ integrin) nor VLA-4 (α_4_β_1_ integrin) were expressed by A20 cells, because they were lacking the β_2_ subunit (CD18) and α_4_ subunit (CD49d), respectively. LFA-1 and VLA-4 mediate lymphocyte stable adhesion and promote its migration through the vessel wall (Pals et al. 2007). The LFA-1 absence prevents the leukocyte emigration to inflammation sites, resulting in a severe immunodeficiency (Drillenburg & Pals 2000; Pals et al. 2007; Page et al. 2012). On the other hand, previous studies have proved that LFA-1 expression provokes cell aggregation limiting its proliferation and its expression is induced by CD21 (complement receptor type 2, CR2, C3d receptor, and mature B-cell marker) (Otsuka et al. 2004a; Otsuka et al. 2004b; Tanimoto et al. 2009; Zhao et al. 2014), whose expression is lost in A20 tumour cells. This suggests that A20 B-cell lymphoma line has jointly selected the loss of CD18 and CD21 expression as tumour proliferative and development advantages, reducing the possible cell interaction and aggregation.

A20 tumour B-cells may have evolved to lose the expression of adhesion molecules and chemokine receptors, which are expressed in normal B cells, altering the traffic of neoplastic B cells to peripheral lymphoid organs with respect to normal B-cells. Therefore, HVEM might not have a role in the dissemination of A20 tumour cells. This could be the explanation for not observing significant differences in the metastatic pattern of tumour spread A20 GFP WT versus A20 GFP HVEM^−/−^.

### Unusual increase of B-cells in the thymus of mice inoculated with A20 B-cell lymphoma line

Normally, the thymus presents a small population of B cells that comprises about 0.1-0.5% of thymocytes, a percentage similar to that of dendritic cells (DCs) (Perera et al. 2013; Lu et al. 2015; Yamano et al. 2015). In our studies, we corroborated that in the thymus of control mice, which had not been inoculated with the different A20 tumour cell lines, the percentage of B cells remained within this normal thymic B-cell range. However surprisingly, in the thymus of mice inoculated with A20 GFP cell line (both WT and HVEM deficient) at intermediate and late stages of B-cell lymphoma progression, appeared significant B-cell percentages. The increase of B-cells in the thymus seems to be promoted by A20 tumour colonization of the thymus. However, the increase in the number of B-cells began even before the detection of A20 tumour in this lymphoid compartment. In homeostatic conditions, the development of thymic B-cells is blocked, in relatively early stages, by the generation of an inhibitory microenvironment (Hashimoto et al. 2002). It may be that thymus microenvironment of mice inoculated with A20 tumour cells had suffered changes, which had promoted the development of B-cells and had also facilitated the implantation of A20 tumour in this hematopoietic structure. However, we would still need to determine the phenotype of this unusual population of B-cells in the thymus, that is, whether they present an endogenous origin (thymic B-cells with IgM^high^ IgD^high^ phenotype), or they have been recruited to the thymus (mature follicular (FO) B-cells with IgM^low^ IgD^high^ phenotype) (Perera et al. 2013).

On the other hand, Thymic B-cells are preferably found in the cortico-medullary region, where they could act as antigen-presenting cells (APCs) eliminating autoreactive CD4 T cells (previously stimulated with the same type of antigen) (Perera et al. 2013; Lu et al. 2015; Perera et al. 2016). Furthermore, studies have demonstrated that thymic B-cells promote regulatory T cells (Treg) development and proliferation (Grey et al. 2014; Lu et al. 2015). For this reason, it would be interesting to study whether this new B-cell population, which appeared in the thymus of mice after A20 tumour implantation, may stimulate Treg cell formation (as well as thymic B-cells) in order to modulate an anti-tumour immune response.

### The loss of HVEM expression in the A20 B-cell lymphoma limits leukocyte infiltration in hepatic tumour nodules at an intermediate stage of tumour development

In the intermediate stage of tumour development, in hepatic tumour nodules, the loss of HVEM expression led to an increase in the number of tumour cells, decreasing TILs proportion. In B and T lymphocytes, the cis HVEM-BTLA complex helps to maintain a functional inactivation state (Croft 2005; Vendel et al. 2009; Shui et al. 2011; Pasero & Olive 2013; Gertner-Dardenne et al. 2013). BTLA knock-out mice exhibit a normal lymphocyte development, but they show T and B cells with a hyperproliferative phenotype after TCR and BCR activation, respectively (Croft 2005; Vendel et al. 2009; Shui et al. 2011; Pasero & Olive 2013; Wu et al. 2007). One possible explanation to the higher presence of A20 GFP HVEM^−/−^ (and consequently a lower TILs proportion) than A20 GFP WT cells in hepatic tumour nodules at day 28 tumour implantation, may be the result of a deficiency in inhibitory cis HVEM-BTLA interaction. It would provoke that the threshold of activation of these A20 tumour cells decreases, making them more susceptible to lack of control in their in vivo proliferation and survival. However, in the late stages of lymphoma development (at day 34 after tumour implantation), there were no differences in tumour cells and TILs number in hepatic tumour nodules formed by A20 GFP HVEM^−/−^ or A20 GFP WT cells. It may occur due to A20 GFP WT cells, in advanced stages of tumour progression, have gotten to evade the inhibitory control cis HVEM-BTLA-mediated, and we were not able to see the proliferative differences concerning the HVEM-deficient A20 tumour cells. Furthermore, A20 GFP WT expresses themselves relatively low levels of HVEM, which would serve as an intrinsic control of their activation and proliferation in the early-intermediate stages of tumour development, but would not serve them in the more advanced stages in which the tumour would have evolved, being able to pass this threshold of inactivation and resemble A20 GFP HVEM^−/−^.

On the other hand, we cannot explain what happens with the loss of HVEM-BTLA interaction in trans, and its relation with the immune cells. Whether HVEM were used by the A20 tumour to evade the action of the immune system, acting as a suppressor ligand for the activation of T cells by binding to BTLA expressed in these cells, its loss would be a disadvantage for the tumour. However, our results demonstrate the opposite because A20 GFP HVEM^−/−^ tumour showed lower recruitment of TILs in hepatic tumour nodules in the intermediate stage of tumour development, concerning A20 GFP WT cells.

Authors have demonstrated that the loss of HVEM expression changes the tumour microenvironment, supporting the development of lymphoma through, for example, the increase in the release of TNF family cytokines (Boice et al. 2016; Verdière et al. 2018). A20 GFP HVEM^−/−^ may be altered the normal function of stromal cells, like Cancer-associated fibroblasts (CAFs) that play an important role promoting tumour growth and progression (Xing et al. 2010; Lakins et al. 2018; Tao et al. 2017). Therefore, it could be very interesting to study the microenvironment changes produced by the loss of HVEM in the A20 B-cell lymphoma, which may contribute to the remolding of extracellular matrix (ECM) preventing or limiting the leukocyte recruitment to the tumour microenvironment (Xing et al. 2010).

## ACKNOWLEDGEMENTS

This work has been supported by grants from the European Regional Development Fund and the Castilla y León regional government (EDU/310/2015, April 10^th^).

## SUPPLEMENTARY FIGURES

**Supplementary Table S1.**
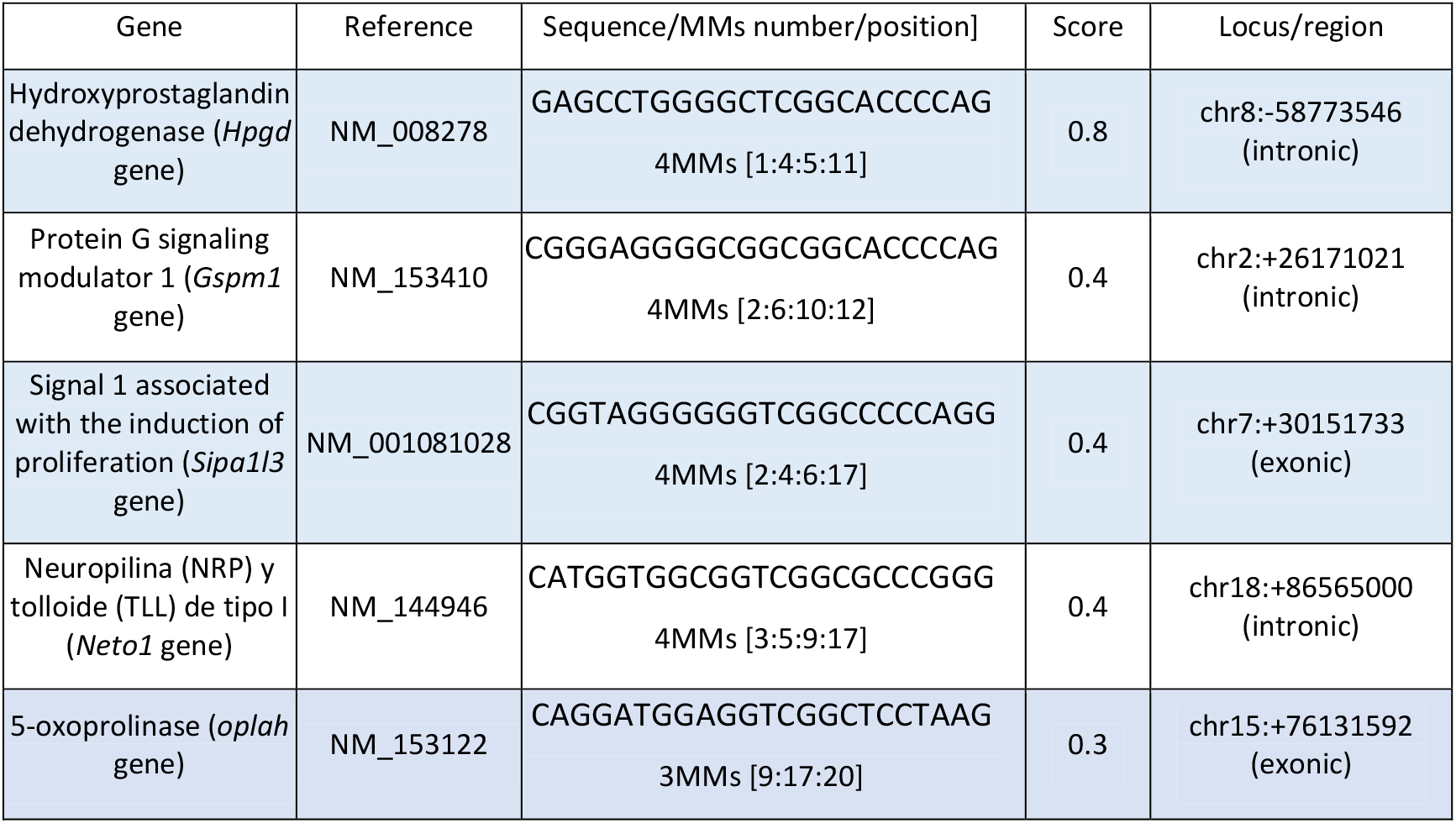
Off-target sequences selected from sgRNA_1. MMs: non-complementary nucleotides or mismatches. The numbers that appear in brackets refer to the positions that MMs occupy in relation to the PAM sequence. For example, *Hpgd* off-target sequence presents the highest off-targeting probability (highest score) because the 4 MMs are located furthest from the PAM sequence (positions 1,4, 5 and 11) with respect to the rest of the off-target sequences analysed.

**Supplementary Figure S1.**
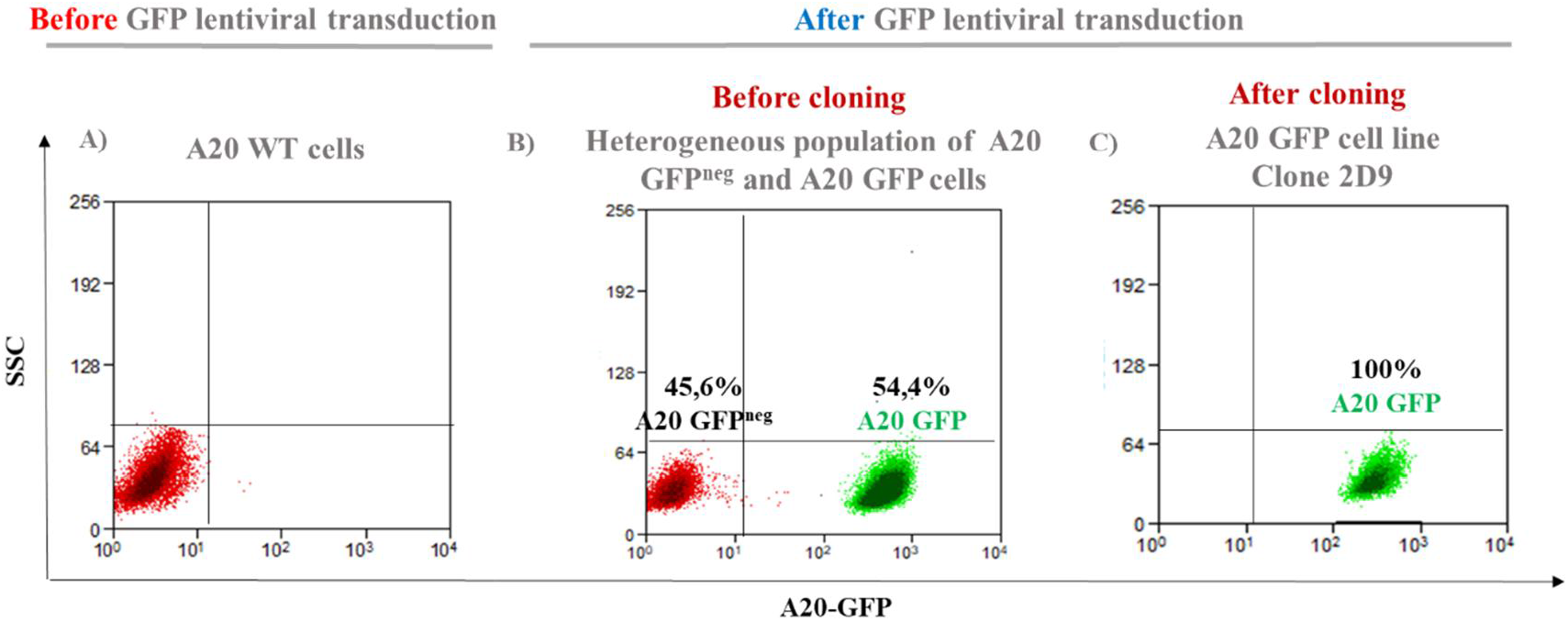
Generation of A20 tumour cell line with GFP stable expression by lentiviral vectors. Representative histogram GFP expression by flow cytometer. (A) All A20 cells were GFP negative before transduction by lentiviral systems encoding GFP. (B) Heterogeneous A20 population as a result of GFP lentiviral transduction, which contained 45.6% A20 GFP negative (GFP^neg^) and 54.4% A20 GFP positive. (C) After cloning by limited dilution of heterogeneous A20 population, we got 100% A20 GFP (clone 2D9) line with GFP stable expression.

**Supplementary Figure S2.**
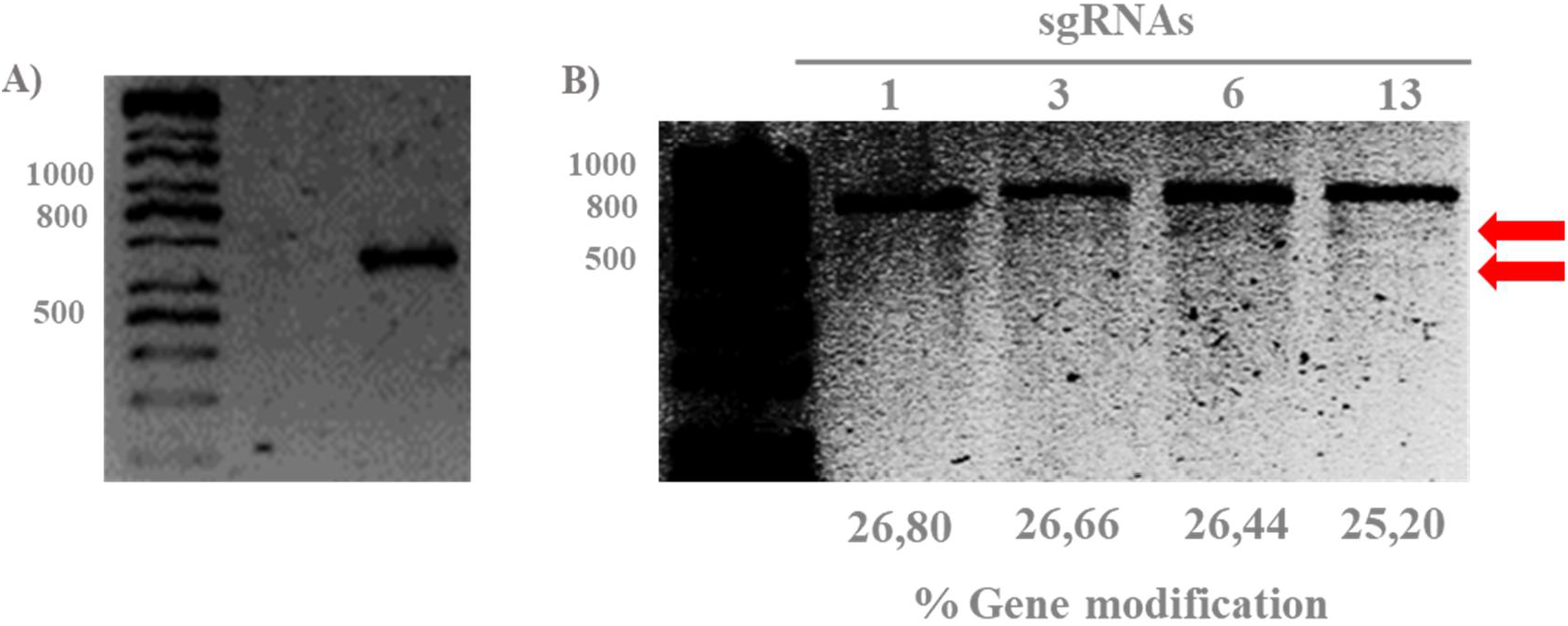
T7 endonuclease I assay shows similar efficiencies of HVEM gene CRISPR-Cas9 modification by different sgRNAs (1, 3, 6 and 13) evaluated. (A) Undigested HVEM band amplified by PCR (cleavage negative control). (B) The resulting digestions of T7/EI assay (red arrows) revealed a similar cleavage pattern to each sgRNA because they presented percentages of HVEM gene modification very similar.

**Supplementary Table S2.** Parameters obtained from the semi-quantification of resulting T7/EI digestion for each sgRNA (1, 3, 6 and 13). We used % Gene modification = 100 × (1− (1-fraction cleaved)^1/2^) formula (Mackay & Segal 2010) (where the fraction cleaved = cleaved product density/ total density (cleaved product + parental peaks)), in order to calculate the cleavage efficiency of each sgRNA, (1, 3, 6 and 13). Density product without cutting was used as a negative control.

| sgRNA | Density product without cutting (Control) | Parental peaks density | Cleaved product density | Total density (cleaved product + parental peaks) | Fraction cleaved | % Gene modification |
| --- | --- | --- | --- | --- | --- | --- |
| (-) | 3723,25 | (-) | (-) | (-) | (-) | (-) |
| 1 | (-) | 3556,29 | 3081,39 | 6637,68 | 0,4642 | 26,80% |
| 3 | (-) | 3324,32 | 2855,64 | 6179,96 | 0,4621 | 26,66% |
| 6 | (-) | 3383,96 | 2869,36 | 6253,32 | 0,4589 | 26,44% |
| 13 | (-) | 3383,36 | 2529,96 | 5913,32 | 0,4405 | 25,20% |

**Supplementary Figure S3.**
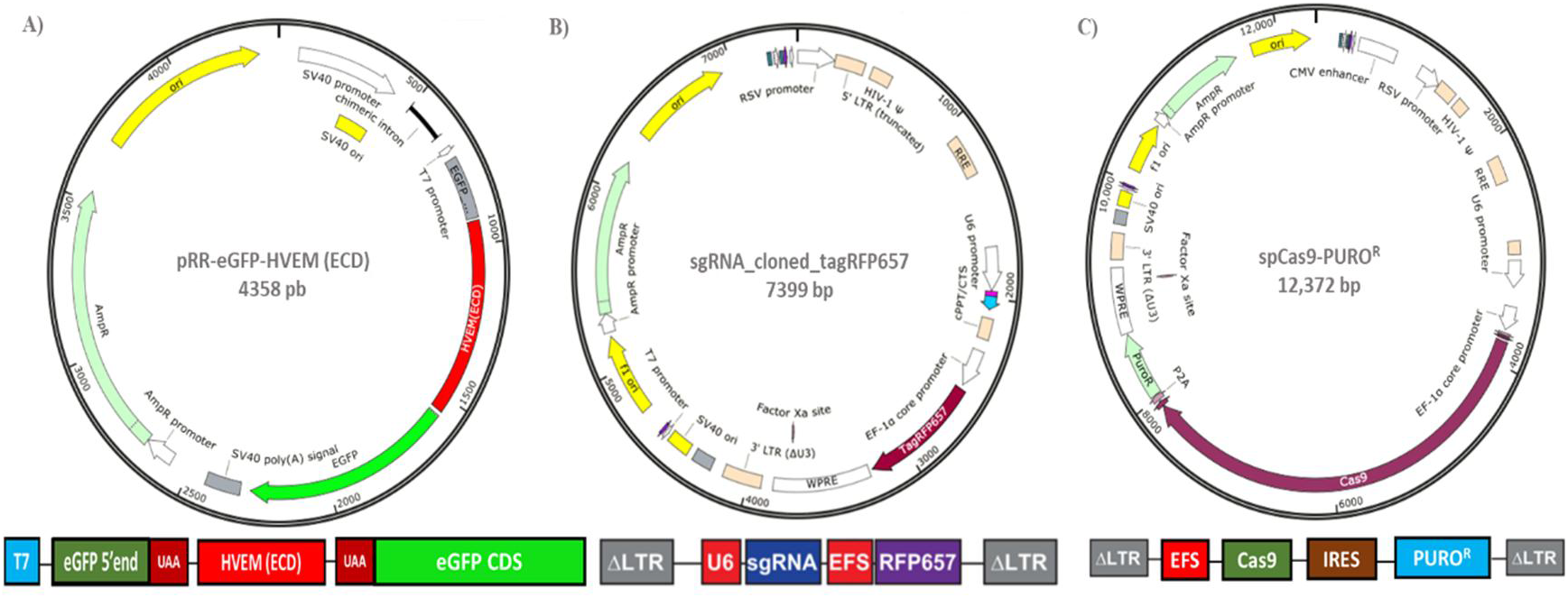
Expression vectors used in order to analyse sgRNA HVEM-cleavage efficiencies by the pRR-eGFP system. (A) pRR-eGFP-HVEM (ECD) vector. HVEM extracellular domain (ECD) (because our CRISPR-Cas9 target site was only of 69 bp corresponding to exon 1) was cloned into pRR-eGFP vector between two sequences coding for eGFP protein (eGFP 5’end: sequence corresponding to the N-terminal end of the eGFP protein and eGFP CDS: rest of the coding sequence (CDS) for the eGFP protein). UAA: stop codon. (B) sgRNA_cloned_tagRFP657. The different designed and selected sgRNAs (1, 3, 6 and 13) were individually cloned into an expression vector that codes for an APC fluorescence protein (tagRFP657) as a reporter gene. (C) spCas9_PURO^R^. Expression vector codes for Cas9 nuclease and puromycin resistant cassette.

**Supplementary Figure S4.**
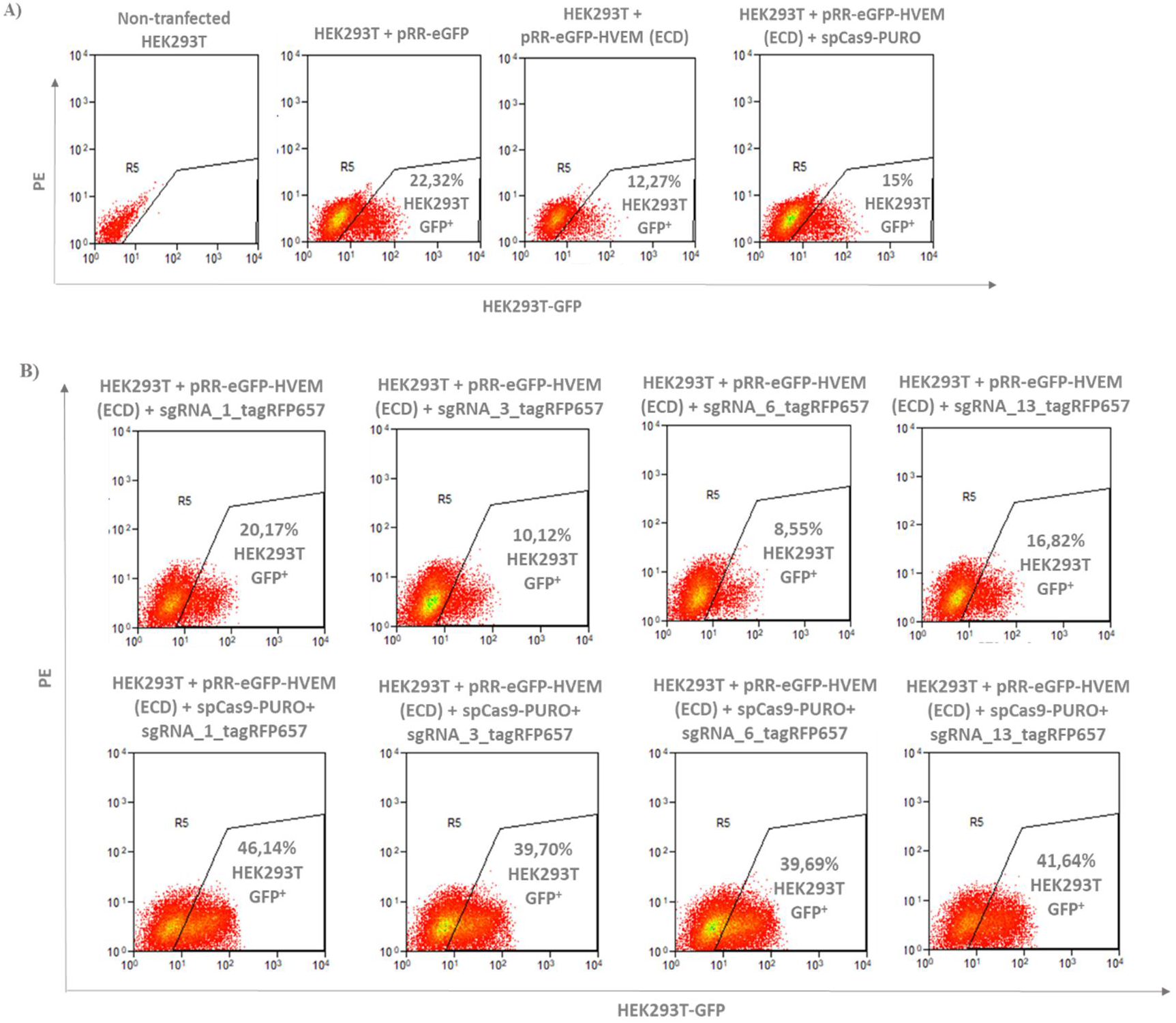
eGFP expression re-establishment after CRISPR-Cas9 action guided by different sgRNAs (1, 3, 6 and 13) exhibits similar HVEM cleavage efficiencies. (A) Non-transfected HEK293T as a negative control. HEK293T cells transfected with empty pRR-eGFP backbone, HVEM (ECD) cloned into pRR-eGFP (pRR-eGFP-HVEM (ECD)), and with pRR-eGFP-HVEM (ECD) and spCas9-PURO (codes for Cas9 nuclease). Empty pRR-eGFP backbone and pRR-eGFP-HVEM (ECD) exhibited basal eGFP auto-fluorescence. (B) In the top line, HEK293T were transfected with pRR-eGFP-HVEM (ECD), and the different sgRNAs without including Cas9 nuclease in order to discount GFP auto-fluorescence. In the bottom line, Cas9 nuclease was included to cleavage HVEM guided by sgRNA and GFP expression re-establishment. The different percentages of GFP positive HEK293T populations were similar and it indicated a similar cleavage pattern to selected sgRNAs.

**Supplementary Figure S5.**
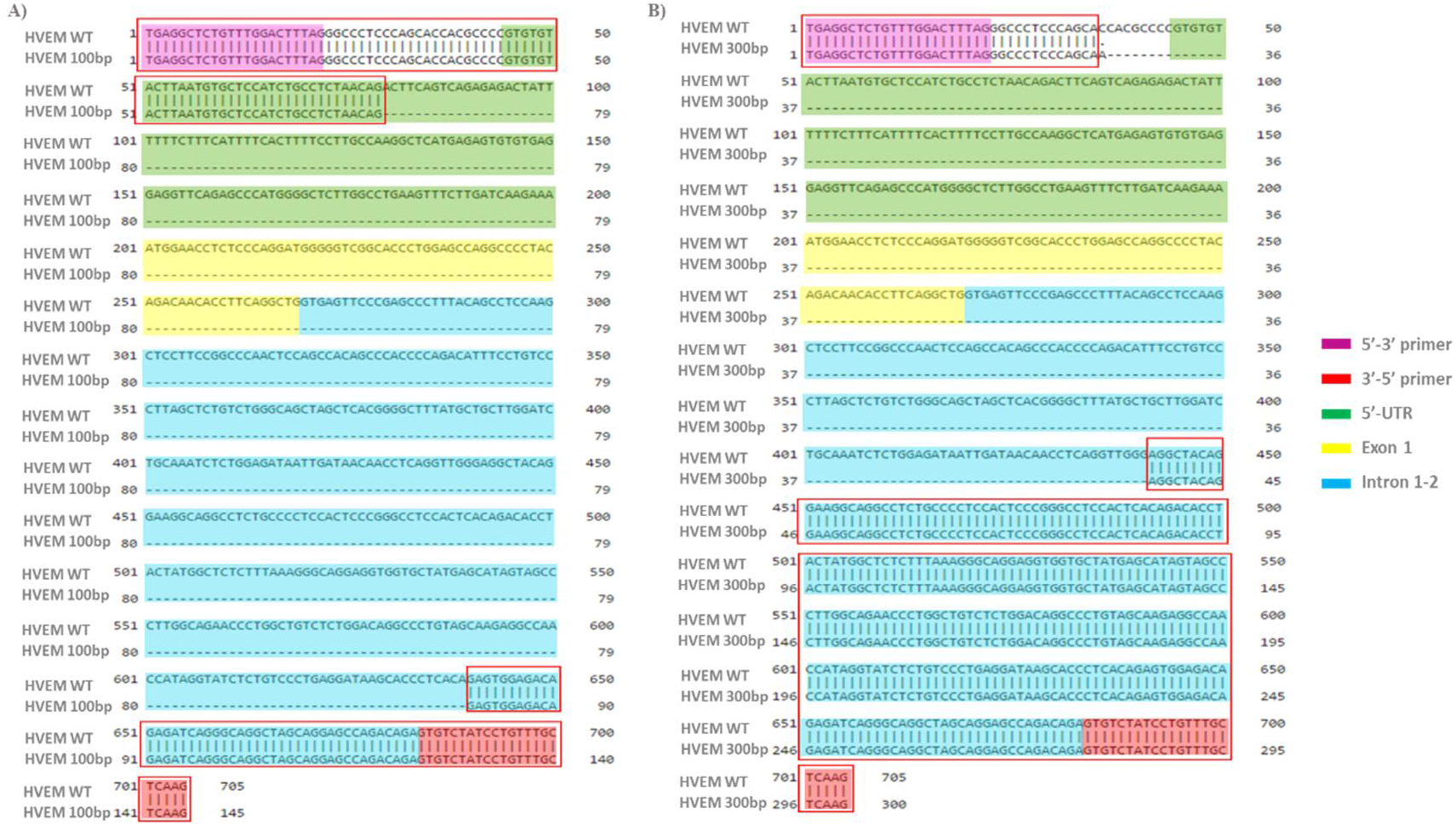
Sequencing analyses reveal two deletions forms of HVEM (100 and 300 bp) by CRISPR-Cas9-sgRNA_1 action. Both deletion mutations (HVEM 100 bp and HVEM 300 bp) were characterized by having lost the complete exon 1 (yellow) and other parts of HVEM sequence had been affected by editing system like 5’-UTR (green) and intron 1-2 (blue) but at a different way. The HVEM 100 bp deletion form (A) conserved a part of 5’-UTR, meanwhile, HVEM 300 bp deletion form (B) had completely lost it. On the other hand, HVEM 100 bp truncated form (A) retained a less intron 1-2 sequence in compare with HVEM 300 bp truncated form (B). Sequences preserved after CRISPR-Cas9-sgRNA editing are included in a red box in both cases. The primers used for PCR amplification of target HVEM region are shaded in purple (5’-3’ primer) and in red (3’-5’ primer).

**Supplementary Table S3.** A20 GFP WT and A20 GFP HVEM^−/−^ cell lines exhibited the same in vitro duplicate rate. In vitro duplicate rate of two cell lines was calculated with material and methods doubling time formula. *Xb*: number of initial cells, 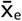: the average of the *Xe* values: count of the number of cells at an incubation time determined (T) (days). The standard deviation (SD) of each value 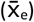 was calculated to determine the degree of statistical significance using the unpaired Student’s t-test method, considering a *p-value* less than 0.05 as statistically significant.

| Cell line | $X_b$ | $\bar{X}_e \pm SD$ (T) | Duplicate rate |
| --- | --- | --- | --- |
| A20 GFP<sup>+</sup> WT (2D9) | $2,5 \times 10^4$ | $4,8533 \times 10^4 \pm 4,6372 \times 10^3$ (T= 1) | 1,04 |
| | | $7,5583 \times 10^4 \pm 9,2680 \times 10^3$ (T= 2) | 1,25 |
| A20 GFP<sup>+</sup> HVEM<sup>-/-</sup> (1F1) | $2,5 \times 10^4$ | $4,7917 \times 10^4 \pm 9,5478 \times 10^3$ (T= 1) | 1,07 |
| | | $7,3667 \times 10^4 \pm 1,6479 \times 10^4$ (T= 2) | 1,28 |

